# Over-expression of Hsp83 in grossly depleted *hsrω* lncRNA background causes synthetic lethality and *l(2)gl* phenocopy in *Drosophila*

**DOI:** 10.1101/420554

**Authors:** Mukulika Ray, Sundaram Acharya, Sakshi Shambhavi, Subhash C. Lakhotia

## Abstract

We examined interactions between Hsp83 and *hsrω* lncRNAs in *hsrω*^*66*^ *Hsp90GFP* homozygotes, which almost completely lack *hsrω* lncRNAs but over-express Hsp83. All *+/+; hsrω*^*66*^ *Hsp90GFP* progeny died before third instar. Rare *Sp/CyO; hsrω*^*66*^ *Hsp90GFP* reached third instar stage but phenocopied *l(2)gl* mutants, dying after prolonged larval life, becoming progressively bulbous and transparent with enlarged brain. Additionally, ventral ganglia were elongated. However, *hsrω*^*66*^ *Hsp90GFP*/*TM6B* heterozygotes, carrying *+/+* or *Sp/CyO* second chromosomes, developed normally. Total RNA sequencing (*+/+, +/+*; *hsrω*^*66*^*/hsrω*^*66*^, *Sp/CyO; hsrω*^*66*^*/hsrω*^*66*^, *+/+; Hsp90GFP/Hsp90GFP*, and *Sp/CyO; hsrω*^*66*^ *Hsp90GFP/hsrω*^*66*^ *Hsp90GFP* late third instar larvae) revealed similar effects on many genes in *hsrω*^*66*^ and *Hsp90GFP* homozygotes. Besides additive effect on many of them, numerous additional genes were affected in *Sp/CyO; hsrω*^*66*^ *Hsp90GFP* larvae, with *l(2)gl* and several genes regulating CNS being highly down-regulated in surviving *Sp/CyO; hsrω*^*66*^ *Hsp90GFP* larvae, but not in *hsrω*^*66*^ or *Hsp90GFP* single mutants. Hsp83 binds at these gene promoters. Several omega speckle associated hnRNPs too may bind with these genes and transcripts. Hsp83-hnRNP interactions are also known. Thus, elevated Hsp83 in altered hnRNP distribution and dynamics, following absence of hsr*ω* lncRNAs and omega speckles, background can severely perturb regulatory circuits with unexpected consequences, including down-regulation of tumor suppressor gene like *l(2)gl*.

## 1. Introduction

The 83 kDa heat shock protein (Hsp83), the *Drosophila* homolog of human Hsp90, is a major member of the Hsp90 family of molecular chaperone proteins (Arya *et al.* 2007; Li *et al.* 2007; Rutherford and Lindquist 1998). Besides being induced by thermal and other cell stress conditions, Hsp83 is ubiquitously expressed during normal development. It is involved in a wide variety of cellular processes like protein chaperoning, chromatin modification through binding with DNA, formation of pre-RISC complex, piwiRNA biogenesis, sno/sn/telomerase RNA accumulation, cell cycle regulation, signaling pathways, transgenerational inheritance and molecular evolution (Bandura *et al.* 2013; Basto *et al.* 2007; Boulon *et al.* 2010; Creugny *et al.* 2018; Cutforth and Rubin 1994; DeFranco and Csermely 2000; Dittmar and Sen 2018; Gangaraju *et al.* 2011; Graf and Enver 2009; Iwasaki *et al.* 2015; Lange *et al.* 2000; Mazaira *et al.* 2016; Miyoshi *et al.* 2010; Olivieri *et al.* 2012; Pratt and Toft 1997; Ruden and Lu 2008; Rutherford *et al.* 2007; Sawarkar *et al.* 2012; Specchia *et al.* 2010; Tariq *et al.* 2009; van der Straten *et al.* 1997; Zhao *et al.* 2008).

An earlier study in our lab (Lakhotia and Ray 1996) showed that heterozygosity for *hsp83* mutant allele in *Drosophila* enhanced the lethality associated with a null condition for the *hsr*□ gene, which produces multiple long non coding RNAs (lncRNA). This indicated possible interaction between Hsp83 and *hsrω* lncRNAs. Like the Hsp83, the *hsr*□ lncRNAs are ubiquitously present in nearly all tissues of *Drosophila melanogaster* at every stage of development and are highly up-regulated under cell stress condition (Lakhotia 2011). Some of the *hsrω* transcripts are cytoplasmic while others are nuclear; the latter are known to associate with diverse hnRNPs and some other proteins to organize the nucleoplasmic omega speckles and thus regulate dynamics of these important regulatory proteins (Lakhotia 2011, 2016; Lakhotia 2017; Piccolo *et al.* 2014; Prasanth *et al.* 2000; Singh and Lakhotia 2015). The Hsp83, although majorly a cytoplasmic protein, is also present in association with perichromatin ribonucleoproteins (Carbajal *et al.* 1990), on many chromosome sites in normal and heat shocked cells (Morcillo *et al.* 1993; Sawarkar *et al.* 2012; Tariq *et al.* 2009). Following heat shock at 37°C, this protein also shows a strong presence at the 93D puff, where the *hsr*□ gene is located (Morcillo *et al.* 1993; Tariq *et al.* 2009).

To further study possible interactions between Hsp83 and *hsrω* transcripts, here we examined effect of over-expression of Hsp83 in a background where the *hsrω* transcripts are nearly absent. We used a transgenic line (Tariq *et al.* 2009) expressing EGFP tagged Hsp83 protein under endogenous *hsp83* promoter to over-express Hsp83. This transgenic line, referred to as *Hsp90GFP* in the following, carries four copies (2 normal and two EGFP-tagged transgenes) of the *hsp83* gene and, therefore, over produces Hsp83 although without any adverse effect on normal development (Tariq *et al.* 2009). The *hsr*□^*66*^ mutant allele shows extremely reduced expression of *hsr*□ transcripts and in homozygous condition causes high degree of lethality so that only about 30% of the *hsr*□ ^*66*^ homozygous progeny survive as weak, short-lived and poorly fertile adults (Johnson *et al.* 2011; Lakhotia *et al.* 2012). Surprisingly, however, when brought together, all of the *+/+; hsr*□ ^*66*^ *Hsp90GFP* homozygous, but not *+/+; hsr*□^*66*^ *Hsp90GFP/TM6B* heterozygous, progeny died at early larval stages. On the other hand, a few *Sp/CyO; hsr*□ ^*66*^ *Hsp90GFP* progeny survived to 3^rd^ instar stage but all of them died after a prolonged third instar life as larvae or, rarely, as pseudopupae. The phenotypes of these larvae, like bloated appearance, deformed imaginal discs, enlarged brain lobes (BL), small endoreplicated organs like salivary and ring glands, gut regions etc, were strongly reminiscent of the phenotypes displayed by the apico-basal polarity *l(2)gl* gene mutant larvae (Aggarwal and King 1969; Farkas and Mechler 2000; Gateff and Mechler 1989; Korochkina *et al.* 1975; Welch 1957). Analysis of the larval transcriptomes of different genotypes revealed that among the many different pathways and genes affected in *hsrω*^*66*^ or *Hsp90GFP* genotypes, several were commonly affected in either of the genotypes. However, the *hsrω*^*66*^ *Hsp90GFP* individuals showed uniquely differential expression of a much large number of genes involved in important processes like cell adhesion, cell fate and proliferation, tissue patterning and differentiation, protein folding and degradation, metamorphosis, microtubule/axoneme formation etc, which together seem to be responsible for the complete lethality of *+/+; hsrω*^*66*^ *Hsp90GFP* homozygotes before the 3^rd^ instar stage. Survival of the few *Sp/CyO; hsrω*^*66*^ *Hsp90GFP* larvae beyond the 2^nd^ instar stage suggests that some genetic factors associated with the *Sp/CyO* chromosomes partially rescued the early damage. However, the surviving *Sp/CyO; hsrω*^*66*^ *Hsp90GFP* larvae displayed greatly reduced levels of transcripts of the apico-basal polarity *l(2)gl* gene and some others that are involved in morphogenesis of brain lobe and ventral ganglia. Since none of these genes were affected in *ω* or *Hsp90GFP* single mutants or in *hsrω Hsp90GFP/+*heterozygotes, interactions between these two gene products appear to be very sensitive to their relative levels. Depletion of *hsrω* transcripts affects intra-cellular dynamics of the diverse hnRNPs (Lakhotia *et al.* 2012;Piccolo *et al.* 2017; Piccolo and Yamaguchi 2017) while Hsp83/Hsp90 is known to bind (Sawarkar *et al.* 2012; Singh and Lakhotia 2015; Tariq *et al.* 2009) with promoters of several highly down-regulated genes in *Sp/CyO; hsrω*^*66*^ *Hsp90GFP* larvae. Hsp90 and a variety of hnRNPs also directly interact and regulate a wide variety of cellular activities (Chen *et al.* 2007; Chi *et al.* 2018; Creugny *et al.* 2018; Ford *et al.* 2002; Grammatikakis *et al.* 1999; Jinwal *et al.*2012; Lackie *et al.* 2017; Youn *et al.* 2018; Zhang *et al.* 2006). Apparently, the altered dynamics of hnRNPs due to the near absence of *hsrω* transcripts in *hsrω*^*66*^ in conjunction with elevated Hsp83 levels severely affects transcripts of many genes, including those like *l(2)gl, kuz, mmp2, SPARC* etc, to generate the unusual phenotype of the rare surviving *Sp/CyO; hsrω*^*66*^ *Hsp90GFP* larvae. These results, besides confirming interactions between *hsrω* transcripts and Hsp83 during development, also highlight that perturbation in interaction between ubiquitously expressed coding and non-coding genes can synergistically result in unexpected but dramatically severe consequences, including initiation of timorous growth by down-regulation of important tumor suppressor genes like *l(2)gl*.

## 2. Materials and Methods

### 2.1 Fly stocks

*1.1* All fly stocks and crosses were maintained on standard agar-maize powder-yeast and sugar food at 24±1°C. The following stocks were used

a) +/+ (Oregon R^+^)

b) *w*^*1118*^; *+/+; hsrω*^*66*^ (Johnson *et al.* 2011)

b) *w*^*1118*^; *Sp/CyO; hsrω*^*66*^

c) *w*^*1118*^; *Sp/CyO; Hsp90GFP* (Tariq *et al.* 2009)

d) *w*^*1118*^; *Sp/CyO; hsrω*^*66*^ *Hsp90GFP* (generated by recombining *hsrω*^*66*^ mutant allele with *Hsp90GFP* transgene insertion)

e) *w*^*1118*^; *+/+; hsrω*^*66*^ *Hsp90GFP* (generated by recombining *hsrω*^*66*^ mutant allele with *Hsp90GFP* transgene insertion)

f) *w*^*1118*^; *Sp/CyO; UAS-hsrωRNAi*^*3*^ *Hsp90GFP* (generated by recombining *UAS-hsrωRNAi*^*3*^ transgene insertion (Mallik and Lakhotia 2009) with *Hsp90GFP* transgene insertion)

g) *w*^*1118*^; *Act5C-GAL4/CyO; +/+*

Appropriate crosses were made, when required, to obtain progeny of the desired type.

The *w*^*1118*^; *Sp/CyO; Hsp90GFP* stock (Tariq *et al.* 2009) was provided by Dr. Renato Paro (ETH Zurich, Basel, Switzerland) and *w*^*1118*^; *+/+; hsrω*^*66*^ (Johnson *et al.* 2011) by Dr. S. W Mckechnie (Monash University, Australia).

### 2.2 Lethality and embryo hatching Assay

For lethality assay, freshly hatched 1st instar larvae of the desired mentioned genotypes were collected during a two hr interval and transferred to food vials containing regular food and reared at 24±1°C. Total numbers of larvae that pupated and subsequently emerged as flies were counted. At least three biological replicates of each experimental condition and/or genotypes were examined.

To find out embryo hatchability, total number of eggs laid during a period of two hours was counted and the numbers of dead and hatched embryos noted for each genotype.

### 2.3 Whole organ immunostaining

Brain complex and salivary glands from actively migrating late third instar larvae of the desired genotypes were dissected out in Poels’ salt solution (Tapadia and Lakhotia 1997), and immediately fixed in freshly prepared 3.7% paraformaldehyde in PBS for 20 min and processed for immunostaining, as described earlier (Prasanth *et al.* 2000), with mouse anti-Wingless (4D4, Developmental Studies Hybridoma Bank, Iowa, 1:50), mouse anti-Dlg (4F3, Developmental Studies Hybridoma Bank, Iowa, 1:50), rabbit anti-PH3 (Phospho-Histone H3 (Ser10), Cell signaling, 1:50), or Anti-Hsp83 (gifted by Dr. Robert Tanguay, 3E6, Morcillo et al 1993; 1:50). Appropriate secondary antibodies conjugated either with Cy3 (1:200, Sigma-Aldrich, India) or Alexa Fluor 633 (1:200; Molecular Probes, USA) or Alexa Fluor 546 (1:200; Molecular Probes, USA) were used to detect the given primary antibody. Chromatin was counterstained with DAPI (4’, 6-diamidino-2-phenylindole dihydrochloride, 1μg/ml). Tissues, mounted in DABCO antifade mountant, were examined under Zeiss LSM Meta 510 confocal microscope using Plan-Apo 40X (1.3-NA) or 63X (1.4-NA) oil immersion objectives. All images were assembled using the Adobe Photoshop software.

The Anti-Hsp83 was also used for western detection of Hsp83 at 1:500 dilution using total proteins samples from wild type, *Sp/CyO; hsrω*^*66*^, *Sp/CyO; Hsp90GFP*, and *Sp/CyO; hsrω*^*66*^ *Hsp90GFP* larvae, as described earlier (Singh and Lakhotia 2015).

### 2.4 RNA isolation and Reverse Transcription-PCR

Total RNA, isolated from healthy wandering third instar larvae of the desired genotypes using Trizol reagent following the manufacturer’s (Sigma-Aldrich, India) recommended, were used for semi-quantitative RT-PCR or qRT-PCR. One μg of RNA, suspended in nuclease-free water and quantified spectrophotometrically, was incubated with 2U of RNase free DNAseI (MBI Fermentas, USA) for 30 min at 37°C to remove any residual DNA. First strand cDNA was synthesized from 1-2 μg of total RNA as described (Mallik and Lakhotia 2009). The cDNAs were subjected to semi quantitative RT-PCR or real time PCR using the desired forward and reverse primer pairs (see Table 1). Real time qPCR was performed using 5µl qPCR Master Mix (SYBR Green, Thermo Scientific), 2 picomol/µl of each primer per reaction in 10 µl of final volume in an ABI 7500 Real time PCR machine. The PCR amplification reactions were carried out in a final volume of 25 μl containing cDNA (50 ng), 25 pM each of the two specified primers, 200 μM of each dNTPs (Sigma Aldrich, USA) and 1.5U of *Taq* DNA Polymerase (Geneaid, Bangalore). 15 μl of the PCR products were loaded on a 2% agarose gel to check for amplification, with a 50bp DNA ladder used as molecular marker.

**Table 1.**
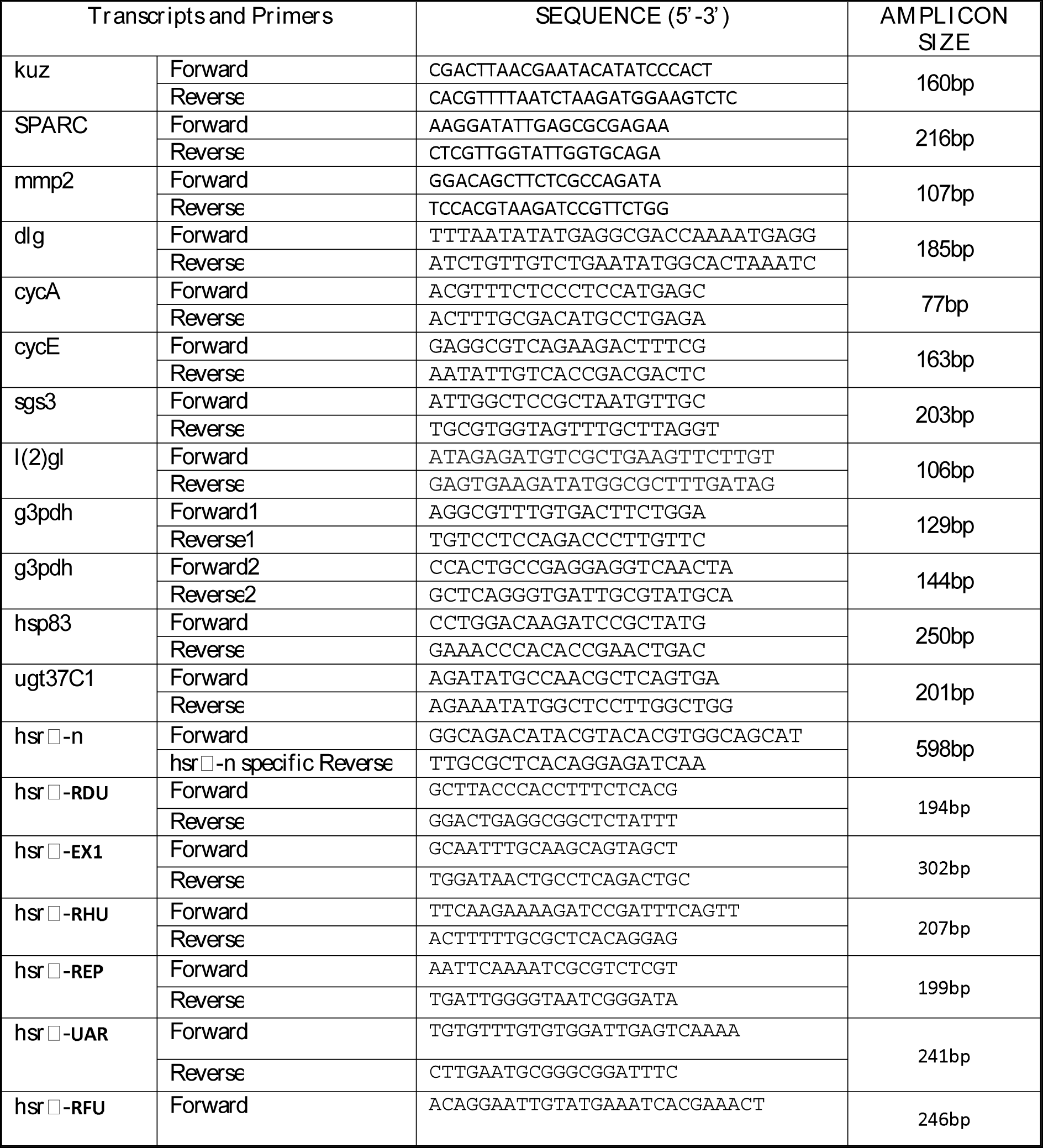
Forward and reverse primers used for qRT-PCR/semi quantitative RT-PCR for quantification of different transcripts

### 2.5 Next Generation RNA sequencing

Total RNA, in duplicate biological replicates, was isolated from *Oregon R*^*+*^ (WT), *Sp/CyO; hsrω*^*66*^, *Sp/CyO; Hsp90GFP, Sp/CyO; hsrω*^*66*^ *Hsp90GFP* wandering third instar larvae (115-117hr after egg hatching) using Trizol (Invitrogen, USA) reagent as per manufacture’s protocol in one set and WT and *+/+; hsrω*^*66*^ in another set. 1µg of each of the isolated RNA samples was subjected to DNAse treatment using 2U of TurboTM DNAse (Ambion, Applied Biosystem) enzyme for 30 min at 37°C. The reaction was stopped using 15mM EDTA followed by incubation at 65°C for 5-7 min and purification on RNAeasy column (Qiagen). The first set of samples (WT, *Sp/CyO; hsrω*^*66*^, *Sp/CyO; Hsp90GFP, Sp/CyO; Hsp90GFP hsrω*^*66*^) were sequenced at the NGS facility at the Centre for Genetic Disorders at Banaras Hindu University while total RNA sequence for the 2nd set (WT and *+/+; hsrω*^*66*^) of samples was provided by Bionivid Technology Private Limited, Bangalore (India). In each case, the biologically duplicate purified RNA samples were processed for preparations of cDNA libraries using the TruSeq Stranded Total RNA Ribo-Zero H/M/R (Illumina) and sequenced on HiSeq-2500 platform (Illumina) using 50bp (first set) or 100bp (2^nd^ set) pair-end reads. For the first set, 8 samples were run per lane, with duplicate samples run across 2 lanes while for the second set, 4 samples were run per lane, with duplicate samples run across 2 lanes. This resulted in a sequencing depth of ~20 million reads in each case. The resulting sequencing FastQ files were mapped to the *Drosophila* genome (Relese dm6) using Tophat with Bowtie. The aligned SAM/BAM file for each was processed for guided transcript assembly using Cufflink 2.1.1 and gene counts were determined. Transcripts from all samples were subjected to Cuffmerge to get final transcriptome assembly across samples. A comparison of the WT RNA sequence outputs from the two sets indicated that they are generally similar. In order to ascertain differential expression of gene transcripts between different samples, Cuffdiff 2.1.1 was used (Trapnell *et al.* 2012). A P-value <0.05 was taken to indicate significantly differentially expressing genes between different groups compared. Genes differentially expressed between experimental and control genotypes were categorized into various GO terms using DAVID bioinformatics Resources 6.8 (Huang *et al.* 2009) (https://david.ncifcrf.gov) for gene ontology search. In order to find out distribution of differentially expressing genes into various groups, Venn diagrams and Heat maps were prepared using the Venny2.1 and ClustVis softwares, respectively (Metsalu and Vilo 2015).

## 3. Results

### 3.1 Synthetic lethality and l(2)gl phenocopy following over expression of Hsp83 in hsr□ RNA depleted condition

The *hsrω*^*66*^ allele is associated with a 1598bp deletion of the upstream regulatory sequences including 9bp downstream of the second transcription start site of the *hsrω* gene and therefore, it has been described as a null allele (Johnson *et al.* 2011). In order to confirm this, we carried out qRT-PCR with total RNAs from *hsrω*^*66*^/*hsrω*^*66*^, *hsrω*^*66*^/*TM6B* and *Act5C-GAL4/CyO; UAS-hsrωRNAi*^*3*^*/TM6B* larvae using several different primer pairs that would amplify transcripts corresponding to different regions of the ~21kb long *hsrω* gene (Fig. 1A and Table 1). Data in Table 1 show that as expected the RD transcript is completely absent in *hsrω*^*66*^ homozygotes while the transcripts corresponding to other parts of the *hsrω* gene are greatly reduced although not completely absent. As shown in Fig. 1A, the exon 1 region is present in all the *hsrω*transcripts, the UBR is common to RH, RB and RF transcripts, repeats and UAR regions are common to RB (and its spliced product RG) and RF transcripts while the FUR is unique to the *hsrω-RF* transcripts. The variable down-regulation of these different transcripts in the *hsrω*^*66*^ homozygotes is due to the variable relative abundance of the different transcripts and to their varying commonality in the multiple transcripts. The present data thus show that the *hsr*□^*66*^ is a very strong hypomorphic allele whose homozygosity results in most of the *hsrω* transcripts being expressed only in trace amounts. The qRT-PCR results with total RNA from *hsrω*^*66*^/*TM6B* heterozygotes showed that all the *hsrω* transcripts, except the RD, were significantly reduced when compared with wild type (Table 1). *Act5C-GAL4* driven ubiquitous expression of the *UAS-hsrωRNAi*^*3*^ transgene, which primarily targets the 280b repeats in the *hsrω* transcripts (Mallik and Lakhotia 2009), also significantly reduced levels of different *hsrω* transcripts, although the degree of reduction was less marked than even in the *hsr*□ ^*66*^*/TM6B* larvae (Table 2).

**Fig. 1.**
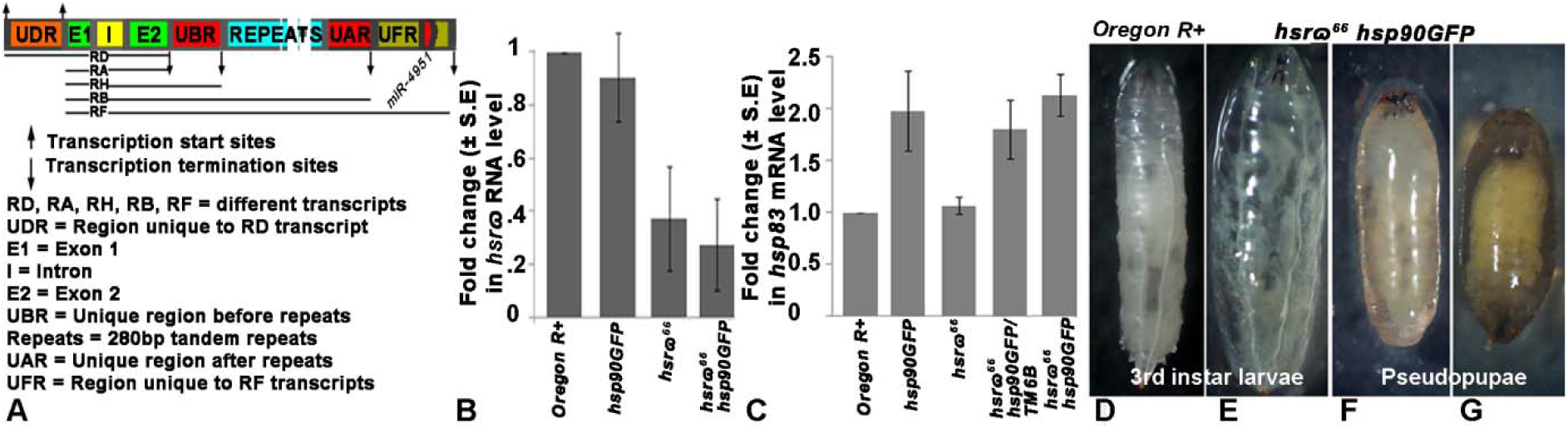
*Sp/CyO; hsr*□^*66*^ *Hsp90GFP* individuals with elevated Hsp83 and depleted *hsr*□ transcripts have prolonged larval life and none form viable pupae. **A** Schematic of the *hsrω* gene (not to scale; for details see http://flybase.org/reports/FBgn0001234) showing its different transcripts and different regions whose transcript levels were examined in Table 2. **B-C** Histograms showing relative fold changes (±S.E, Y-axis) in levels of *hsr*□ (**B**) and *hsp83* transcripts (**C**) following semi-quantitative RT-PCR in different genotypes (X-axis). **D-G** Images of wild type late third instar larva (**D**), *hsr*□^*66*^ *Hsp90GFP* larva (**E**) and *hsr*□ ^*66*^ *Hsp90GFP* pseudopupae (**F, G**).

**Table 2.** Different *hsrω* transcripts are nearly absent in *hsr ω*^*66*^ homozygotes

| Genotype | Fold changes in transcripts corresponding to different regions of the *hsrω* gene* |  |  |  |  |  |
| --- | --- | --- | --- | --- | --- | --- |
|  | RDU | Exon1 | UBR | Tandem Repeats | UAR | FUR |
| WT | 1.0 | 1.0 | 1.0 | 1.0 | 1.0 | 1.0 |
| *hsrω<sup>66</sup>/hsrω<sup>66</sup>* | - | -60.18 | -1371.54 | -2879.42 | -109.54 | -1085.34 |
| *hsrω<sup>66</sup>/TM6B* | 1.77 | -4.99 | -11.32 | -45.36 | -6.08 | -15.11 |
| *Act5C-GAL4/CyO; UAS-hsrωRNAi<sup>3</sup>/TM6B* | -7.49 | -1.29 | -7.74 | -2.888 | -2.57 | -2.24 |

\* Different regions (RDU, Exon1, UBR, Tandem Repeats, UAR and FUR) of the *hsr $\omega$* gene are schematically shown in Fig. 1A.

We also compared levels of *hsrω* transcripts in *Oregon R*^*+*^ (wild type WT), *Hsp90GFP/Hsp90GFP, hsrω*^*66*^/*hsrω* ^*66*^ and, *hsrω* ^*66*^ *Hsp90GFP*/*hsrω* ^*66*^ *Hsp90GFP* by semi-quantitative RT-PCR using primer pairs corresponding to exon 1 so that all the *hsrω* transcripts get amplified (Fig. 1B). Levels of *hsrω* transcripts in WT and *Hsp90GFP* homozygotes were comparable but greatly reduced in *hsrω*^*66*^ and *hsrω* ^*66*^ *Hsp90GFP* homozygotes as expected.

Levels of *hsp83* transcripts as detected by semi-quantitative RT-PCR in *Oregon R*^*+*^ (wild type WT) and *hsr*□^*66*^ homozygotes were comparable (Fig. 1C). The *Hsp90GFP* homozygous larvae, with four copies of the *hsp83* gene (Tariq *et al.* 2009) showed nearly double the levels of *hsp83* transcripts compared to WT larvae, but with no change in levels of *hsr*□ transcripts (Fig. 1B, C). The *hsp83* transcript level in *Hsp90GFP/TM6B* larvae, with three copies of *hsp83* gene, was about 1.5 fold greater than that in wild type larvae (Fig. 1B). Levels of *hsr*□ and *hsp83* transcripts in *hsr*□^*66*^ *Hsp90GFP* homozygous larvae were comparable to those in *hsr* ^*66*^ or *Hsp90GFP* homozygotes, respectively (Fig. 1B, C). As reported earlier (Tariq *et al.* 2009), our western-blot analysis (not shown) confirmed that the Hsp83 protein levels in *Hsp90GFP* and *hsr*□^*66*^ *Hsp90GFP* homozygous larvae were greatly elevated than in WT or *hsr*□ larvae. Thus the *hsr*□^*66*^ *Hsp90GFP* larvae had over-abundance of Hsp83 but gross depletion of the *hsr*□ lncRNAs.

In agreement with earlier report (Tariq *et al.* 2009), the viability of *Hsp90GFP* homozygous larvae under normal laboratory rearing conditions was comparable to that of wild type, with no significant larval or pupal death. As reported earlier (Johnson *et al.* 2011; Lakhotia *et al.* 2012), nearly all of the *hsr*□^*66*^ larvae pupated but there was embryonic as well as late pupal lethality so that less than 30% of the *hsr*□^*66*^ homozygous eggs hatched as adults. During the course of our study, we noticed that out of nearly 600 eggs laid by *+/+*; *hsr*□^*66*^ *Hsp90GFP/TM6B* parents, no non-tubby larvae (*+/+*; *hsr*□^*66*^ *Hsp90GFP/hsr*□ ^*66*^ *Hsp90GFP*) were visible beyond first or second instar stage. However, in the progeny of *Sp*/*CyO*; *hsr*□^*66*^ *Hsp90GFP/TM6B* parents, a very small proportion (~5% of 600 eggs examined) of *Sp*/*CyO*; *hsr*□ ^*66*^ *Hsp90GFP* (non-tubby) larvae continued to the third instar stage (Tables 3A, 4). In order to examine if high lethality also occurred during embryonic development in *hsr*□^*66*^ *Hsp90GFP* homozygous embryos, the hatchability of eggs laid by +/+; *hsr*□^*66*^*/TM6B* and *+/+; hsrω*^*66*^ *Hsp90GFP/TM6B* parents and those laid by *Sp/CyO*; *hsr*□^*66*^*/TM6B*, and *Sp/CyO; hsrω*^*66*^ *Hsp90GFP/TM6B* parents was compared. No difference in hatchability of embryos between any of the pairs was found (Table 3B). Together, these results indicate that the viability of *hsrω*^*66*^ *Hsp90GFP* homozygous embryos is comparable to that of *hsrω*^*66*^ homozygous embryos, irrespective of the chromosome 2 constitution (*+/+* or *Sp/CyO*) but the +/+; *hsrω* ^*66*^ *Hsp90GFP/hsrω* ^*66*^ *Hsp90GFP* individuals showed 100% lethality at first or second instar stage while a small proportion of *Sp/CyO; hsr ω*^*66*^ *Hsp90GFP/hsrω*^*66*^ *Hsp90GFP* larvae continued to the 3rd instar stage.

**Table 3.** Differential survival of *+/+; hsrω* ^*66*^ *Hsp90GFP* and *Sp/CyO; hsrω* ^*66*^ *Hsp90GFP* embryos and larvae

| **A. Survival of +/+; *hsrω<sup>66</sup>* *Hsp90GFP/TM6B* and *Sp/CyO; hsrω<sup>66</sup>* *Hsp90GFP/TM6B* to third instar stage** |  |  |
| --- | --- | --- |
| Parental Genotype | No of Replicates | Mean % (± S.D.) non tubby third instar larvae |
| +/+; *hsrω<sup>66</sup>* *Hsp90GFP/TM6B* | 4 (150 first instar larvae each) | 0±0 |
| *Sp/CyO; hsrω<sup>66</sup>* *Hsp90GFP/TM6B* | 4 (150 first instar larvae each) | 4.5±1.46 |
| **B. Survival of +/+; *hsrω<sup>66</sup>* *Hsp90GFP/TM6B* and *Sp/CyO; hsrω<sup>66</sup>* *Hsp90GFP/TM6B* embryos\*** |  |  |
| Genotype of parents | Mean % (± S.D.) dead embryos | Mean % (± S.D.) first instar larvae |
| +/+; *hsrω<sup>66</sup>/TM6B* | 35.64±1.97 | 64.36±1.97 |
| +/+; *hsrω<sup>66</sup>* *Hsp90GFP/TM6B* | 38.41±2.93 | 61.59±2.93 |
| *Sp/CyO; hsrω<sup>66</sup>/TM6B* | 64.60±4.32 | 35.40±4.32 |
| *Sp/CyO; hsrω<sup>66</sup>* *Hsp90GFP/TM6B* | 64.65±3.79 | 35.35±3.79 |

\* About 1000 eggs from 3 replicates with ~350 eggs each were examined for each genotype.

We compared the time required to complete larval and pupal development at 24±1°C in WT, *Hsp90GFP, hsr*□^*66*^, *hsr*□ ^*66*^ *Hsp90GFP/TM6B* (heterozygous for *hsr*□ ^*66*^ and *Hsp90GFP*), and *Sp/CyO; hsr*□^*66*^ *Hsp90GFP/hsr*□ ^*66*^ *Hsp90GFP* genotypes. As the data in Table 4 show, the *Hsp90GFP, hsr*□^*66*^ and *Sp/CyO; hsr*□ ^*66*^ *Hsp90GFP/TM6B* larvae homozygous larvae developed a little slower. In agreement with earlier reports (Johnson *et al.* 2011; Lakhotia *et al.* 2012), *hsr*□^*66*^ showed 15-30% pupal, but no larval lethality while *Hsp90GFP* did not show any larval or pupal lethality. However, none of the *+/+; hsr*□^*66*^ *Hsp90GFP/hsr*□ ^*66*^ *Hsp90GFP* individuals reached the third instar stage, but rare *Sp/CyO; hsr*□^*66*^ *Hsp90GFP/hsr*□ ^*66*^ *Hsp90GFP* larvae (52 out of 580 eggs laid by *Sp/CyO; hsr*□ ^*66*^ *Hsp90GFP/TM6B*) continued as larvae for 16-23 days after hatching (Table 4). These larvae phenocopied *l(2)gl* mutant larvae in their progressively bulbous and transparent appearance (Fig. 1D) and multi-organs defects (see below). Most of them died as larvae. Less than 1% of the delayed larvae formed pseudopupae before dying (Fig. 1E, F). No flies of this genotype ever emerged.

**Table 4.** *+/+; hsr*□^*66*^ *Hsp90GFP/hsr*□^*66*^ *Hsp90GFP* larvae show synthetic lethality and die before 3^rd^ instar stage while the few surviving *Sp/CyO; hsr*□^*66*^ *Hsp90GFP/hsr*□^*66*^ *Hsp90GFP* larvae phenocopy *l(2)gl* mutant larvae

| Genotype | Number of 1<sup>st</sup> instar larvae | Pupation | Adult hatching | Larval Lethality | Pupal Lethality |
| --- | --- | --- | --- | --- | --- |
| *Oregon R+* | 305 | Complete on Day 5 | Complete on Day 10 | Nil | Nil |
| *Hsp90GFP* | 157 | Day 6 to 8 | Day 11 to 12 | Nil | Nil |
| *hsr□<sup>66</sup>* | 103 | Day 5 to 7 | Day 10 to 11 | Nil | 15-20% |
| *Sp/CyO; hsr□<sup>66</sup>* *Hsp90GFP/TM6B* | 155 | Day 5 to 6 | Day 10 to 11 | Nil | Nil |
| +/+; *hsr□<sup>66</sup>* *Hsp90GFP*/*hsr□<sup>66</sup>* *Hsp90GFP* | 600 | None | None | 100% by 1<sup>st</sup>/2<sup>nd</sup> instar stage | Not applicable |
| *Sp/CyO; hsr□<sup>66</sup>* *Hsp90GFP*/*hsr□<sup>66</sup>* *Hsp90GFP* | 52 | None | None | ~99% die between 16-23 days after hatching | <1% die as pseudopupae |

All the *+/+*; *hsr*□^*66*^ *Hsp90GFP/TM6B* and *Act5C-GAL4/CyO; UAS-hsrωRNAi*^*3*^ *Hsp90GFP/TM6B* larvae developed normally without any indication of the phenotypes exhibited by the *Sp/CyO; hsrω*^*66*^ *Hsp90GFP/hsrω* ^*66*^ *Hsp90GFP*.

In subsequent studies, therefore, we examined the rare surviving third instar *Sp/CyO; hsr* □^*66*^ *Hsp90GFP/hsr*□^*66*^ *Hsp90GFP* larvae to understand the reasons for their unusual larval phenotypes. For convenience sake, we refer the *Sp/CyO; hsr*□^*66*^ *Hsp90GFP/hsr*□ ^*66*^ *Hsp90GFP* and *Sp/CyO; Hsp90GFP* genotypes as *hsr*□^*66*^ *Hsp90GFP* and *Hsp90GFP* respectively in the following. The second chromosome constitution is mentioned only when it is not *Sp/CyO*. The *hsr*□^*66*^ genotype indicates that the 2nd chromosomes are wild type but when *Sp/CyO* chromosomes are present in combination with *hsr*□^*66*^ allele on chromosome 3, the genotype is indicated as *Sp/CyO; hsr*□^*66*^.

### 3.2 The rare surviving hsr□^66^ Hsp90GFP third instar larvae show multi-organ defects

Examination of internal organs of the rare *hsr*□^*66*^ *Hsp90GFP* third instar larvae at different stages of their prolonged life revealed developmental defects in nearly every organ, many of which were similar to those seen in *l(2)gl* mutant larvae (Betschinger *et al.* 2003; Farkas and Mechler 2000; Gateff and Mechler 1989; Ohshiro *et al.* 2000; Roy and Lakhotia 1991). The ring gland, which is very conspicuous in *Hsp90GFP/TM6B* late third instar larvae, because of the very high expression of Hsp90GFP (Fig. 2A), was much smaller in *hsr*□^*66*^ *Hsp90GFP* homozygous larvae, even in those which were two days older than the *hsr*□^*66*^ *Hsp90GFP/TM6B* late 3rd instar larvae (Fig. 2B). With advancing age, the ring gland progressively lost its tissue organization in *hsr*□^*66*^ *Hsp90GFP* larvae (Fig. 2C) so that by the 19^th^ day it appeared as an amorphic tissue lump (Fig. 2D). Similar loss of tissue organization was seen in *hsr* □^*66*^ *Hsp90GFP* larval imaginal discs so that their individual identity and the patterned expression of Wingless were lost even by day 6 (Fig. 2E). Likewise, the characteristic distribution patterns of Dlg, a septate junction marker (Woods et al 1996), seen in wild type or *hsr*□^*66*^ (Fig. 2F) or *Hsp90GFP* (Fig. 2G) imaginal discs were also completely lost in *hsr*□^*66*^ *Hsp90GFP* larvae (Fig. 2H-I). The foregut, especially the proventriculus and gastric caecae, were short and stumpy in *hsr*□^*66*^ *Hsp90GFP* compared to WT or *hsr*□^*66*^ or *Hsp90GFP* larvae (Fig. 2J, K). Like the gut, the salivary glands (SG) were also smaller in *hsr*□^*66*^ *Hsp90GFP* larvae than in wild type with fat body cells more globular and sparser (Fig. 2L, M). The endoreplicating cells and their nuclei in SG of 5 day old *hsr*□^*66*^ *Hsp90GFP* larvae were much smaller than in wild type or *hsr*□^*66*^ *Hsp90GFP/TM6B* heterozygous larvae of similar age; their nuclear size showed only a marginal increase during the extended larval life (Fig. 3A-F). It is interesting that the total number of endoreplicating cells in SG of *Hsp90GFP* homozygous larvae was about 10% higher than in wild type or *hsr*□^*66*^; this small but statistically significant increase in total number of cells was also seen in *hsr*□^*66*^ *Hsp90GFP* homozygous larvae (Fig. 3G)

**Fig. 2.**
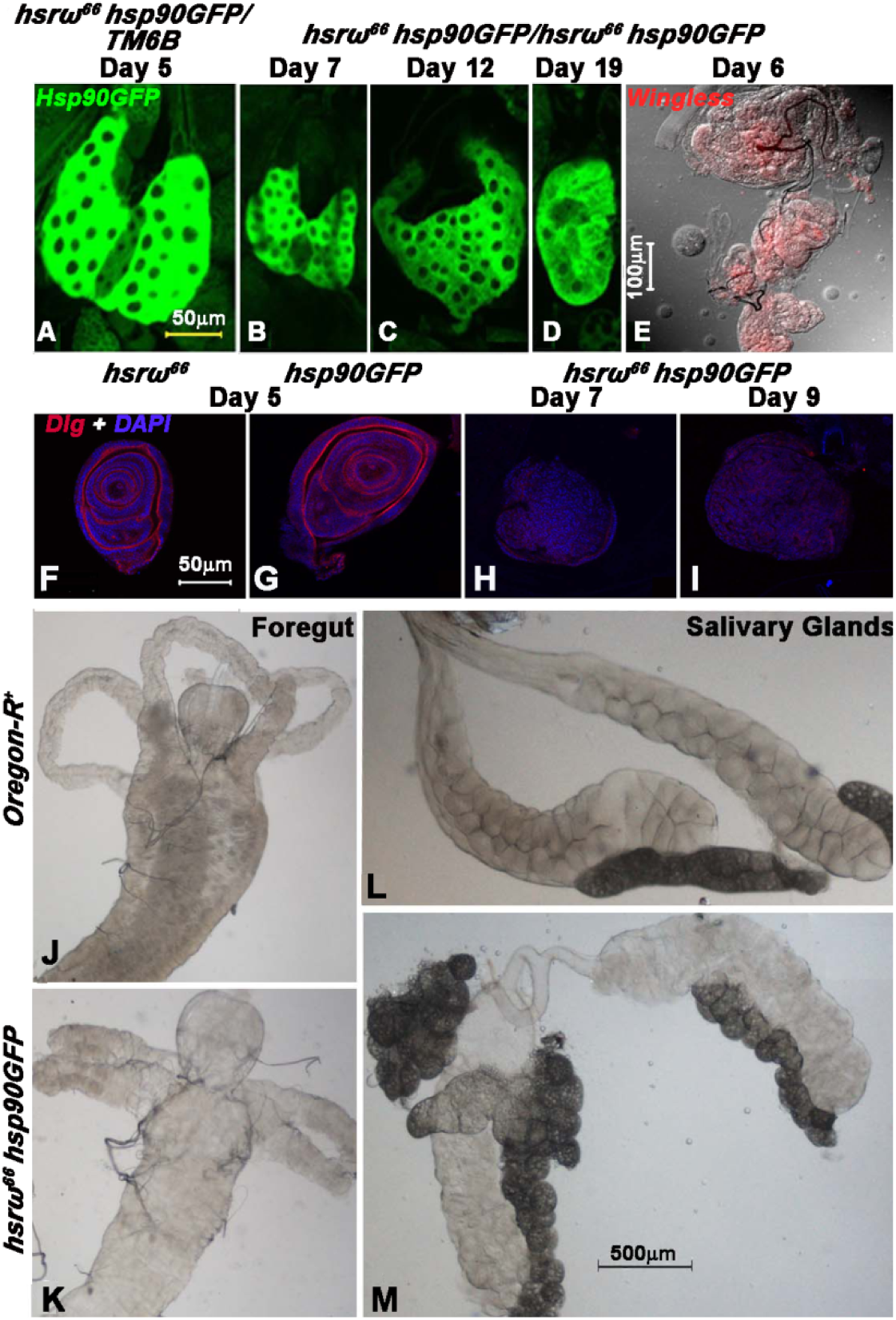
All organs in *hsr*□ ^*66*^ *Hsp90GFP* larvae show defects with endoreplicating organs being smaller in size and mitotically active tissues being disorganized. **A-D** Confocal images showing expression of Hsp90GFP (green,) in ring glands of different larval ages (top of each panel) in *hsr*□^*66*^ *Hsp90GFP/TM6B* (**A**) and *hsr* ^*66*^ *Hsp90GFP* homozygous larvae (**B-D**). **E** Confocal and DIC image of imaginal discs of 6 day old *hsr*□^*66*^ *Hsp90GFP* homozygous larva immunostained with Wingless antibody (red). **F-I** Imaginal discs of *hsr*□^*66*^ (**F**), *Hsp90GFP* (**G**), *hsr*□^*66*^ *Hsp90GFP/TM6B* (**H**-**I**) larvae (larval age on top of the panels). **J-M** DIC images of foregut (**J, K**) and salivary glands (**L, M**) of 5 day old wild type (**J, L**) and *hsr*□^*66*^ *Hsp90GFP* homozygous larvae (**K, M**). Scale bar in **A** applies to **A-D**, that in **F** to **F**-**I**, and that in **M** to **J-M**.

**Fig. 3.**
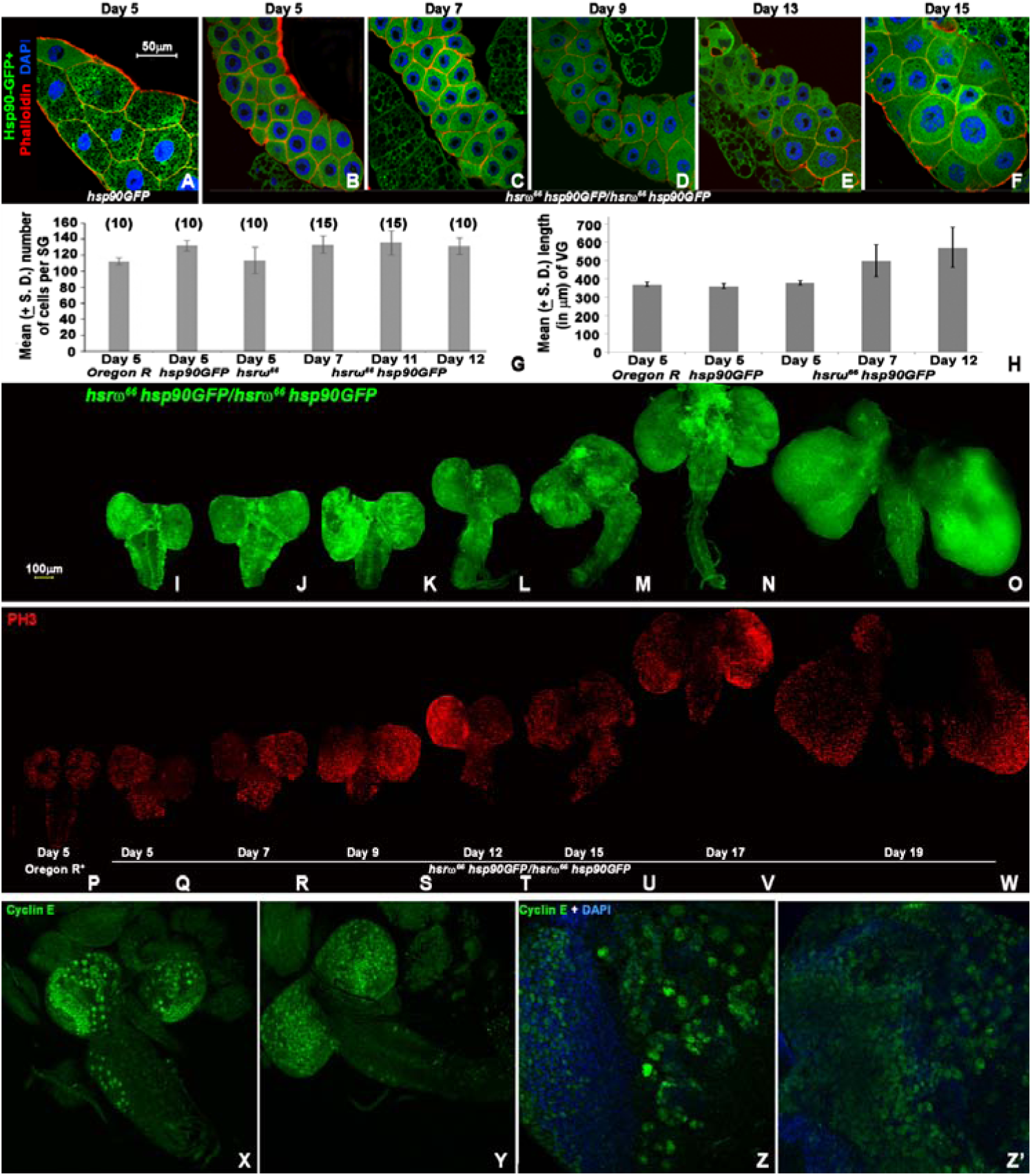
*hsr*□^*66*^ *Hsp90GFP* larvae show small SG throughout the prolonged larval life but progressive increase in sizes of BL and VG. **a-f** Confocal optical sections of salivary glands (SG) of 5 day old *Hsp90GFP* larva (**A**) and *hsr*□^*66*^ *Hsp90GFP* larvae (**B-F**) at different days noted at top of each panel expressing Hsp83GFP (green); cell boundaries are marked by Phalloidin staining for F-actin (red) while nuclei are stained with DAPI (blue). Scale bar in **A** applies to **A-F**. **G** Histogram of mean number (Y-axis) of cells per SG lobe in larvae of different genotypes on different days (X-axis); numbers in parentheses above each bar indicate the number of larvae examined for the given genotype. **H** Histogram showing mean length (Y-axis) of ventral ganglia (VG) in larvae of different genotypes on different days (X-axis). Number of VG measured for each genotype is 15. **I-O** Confocal projection images showing BL and VG of *hsr*□^*66*^ *Hsp90GFP* on different days (noted below each panel) of larval life expressing Hsp83GFP (green). **P-W** Confocal projection images of BL and VG of *Oregon R*^*+*^ (**P**) and *hsr*□^*66*^ *Hsp90GFP* larvae (**Q-W**) of different ages, noted below each panel, showing Phospho-histone 3 (PH3, red) staining; the images in **P-W** are of same BL and VG shown in **I-O** respectively. Scale bar in **G** represents 100µm and applies to **G-W**. X-Z and **Z’** Confocal projection images of whole brain (**X-Y**) and higher magnification images of single brain lobe (**Z-Z’**) of 5 day old wild type (**X, Z**) and *hsr*□^*66*^ *Hsp90GFP* (**Y, Z’**) showing CycE positive cells (green). Blue in **Z** and **Z’** show DAPI positive nuclei.

The most striking feature of the *hsr*□^*66*^ *Hsp90GFP* larvae during their prolonged life was a progressive increase in size of the brain lobes (BL) and the ventral ganglia (VG) (Fig. 3H-O). The length of the VG and size of the BL varied independently in different *hsr*□^*66*^ *Hsp90GFP* homozygous larvae (Fig. 3H-O) so that in some cases, BL were much larger while in others the VG attained a much greater length (e.g., compare Fig. 3N and O). It is notable that 5 day old wild type and *hsr*□^*66*^ *Hsp90GFP* homozygous larvae showed similar sizes of BL and VG (Fig. 3H, P and Q). Immunostaining with mitotic cell marker phosphohistone-3 (PH3) revealed increased staining for PH3 in the BL and at the base of the VG even on day 5 of *hsr*□^*66*^ *Hsp90GFP* larvae, when compared with that in wild type (Fig. 3P, Q). Increased PH3 staining at the basal region of VG during the prolonged larval life (Fig. 3P-W) indicated that active mitosis at the basal region might be responsible for the increase in VG length in *hsr*□^*66*^ *Hsp90GFP* larvae. Immunostaining for Cyclin E (CycE), which is expressed only in undifferentiated neural cells (Bello *et al.* 2008; Berger *et al.* 2010), revealed that compared to wild type (Fig. 3X), the CycE positive cells were much more frequent in BL of *hsrω*^*66*^ *Hsp90GFP* larvae (Fig. 3Y), but the characteristic patterns of the CycE positive cells in the optic lobes including the many large sized CycE positive cells present in WT larvae (Fig. 3Z) were missing in the *hsrω*^*66*^ *Hsp90GFP* larval BL (Fig. 3Z’). This suggests that the stereotyped pattern of divisions of neuroblasts and the subsequent differentiation of their progeny cells (Chia *et al.* 2008; Homem and Knoblich 2012; Lin and Schagat 1997) were absent in BL of *hsrω*^*66*^ *Hsp90GFP*. The CycE staining in *hsrω*^*66*^ or *Hsp90GFP* larval BL was generally similar to that in WT (not shown).

### 3.3 Over expression of Hsp83 in hsr□ RNA depleted background results in down-regulation of large sets of genes involved in cell/tissue division, growth and differentiation

In view of the widespread and severe defects displayed by *hsr*□^*66*^ *Hsp90GFP* larvae, we examined whole transcriptome of 115-117hr (after egg hatching) old *hsr*□^*66*^ *Hsp90GFP* larvae and compared with those of same age wild type (Oregon R), *hsr*□^*66*^, *Sp/CyO; hsr*□^*66*^ and *Hsp90GFP* larvae. We found that 128 and 154 genes were significantly down-regulated in *hsr*□^*66*^ and *Hsp90GFP* larvae, respectively, when compared with similar age wild type larvae (Fig. 4A), with ~35% (45 genes) being common in *hsr*□^*66*^ and *Hsp90GFP* larvae (Fig. 4A and 5A). Similarly, 309 and 232 genes were significantly up-regulated, compared to similar age wild type larvae, in *hsr*□^*66*^ and *Hsp90GFP* larvae, respectively. In this case also ~37% (113) were commonly up-regulated in both these genotypes (Fig. 4B and 5B). Such common effects on a significant proportion of genes in *hsr*□^*66*^ and *Hsp90GFP* genotypes suggest that *hsr*□ transcripts and Hsp83 may be involved in several common pathways. In agreement with this, the heat maps presented in Fig. 5, show that many of the genes that were commonly affected in *hsr*□^*66*^ and *Hsp90GFP* genotypes showed an additive effect in *hsr*□^*66*^ *Hsp90GFP* double mutant larvae.

**Fig. 4.**
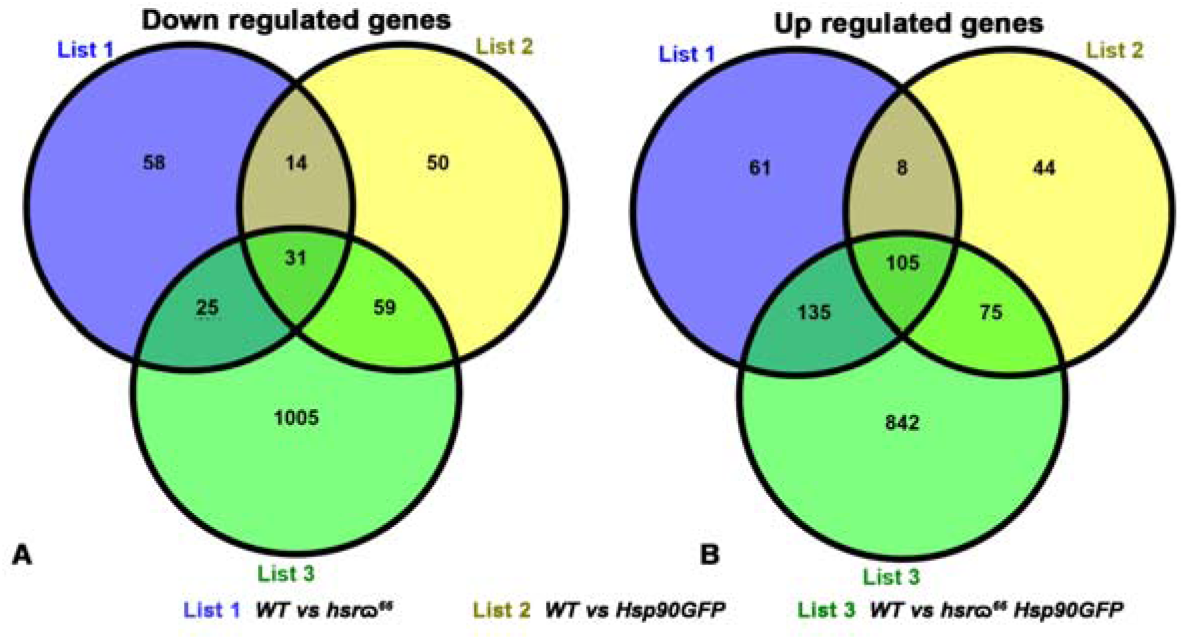
Large numbers of genes show differential expression in 5 day old *hsr*□^*66*^ *Hsp90GFP* larvae compared to wild type, *hsr*□^*66*^ or *Hsp90GFP* larvae of same age. Venn diagrams showing numbers of genes that are significantly down- (**A**) or up-regulated (**B**) in different genotypes when compared with wild type (WT).

**Fig. 5.**
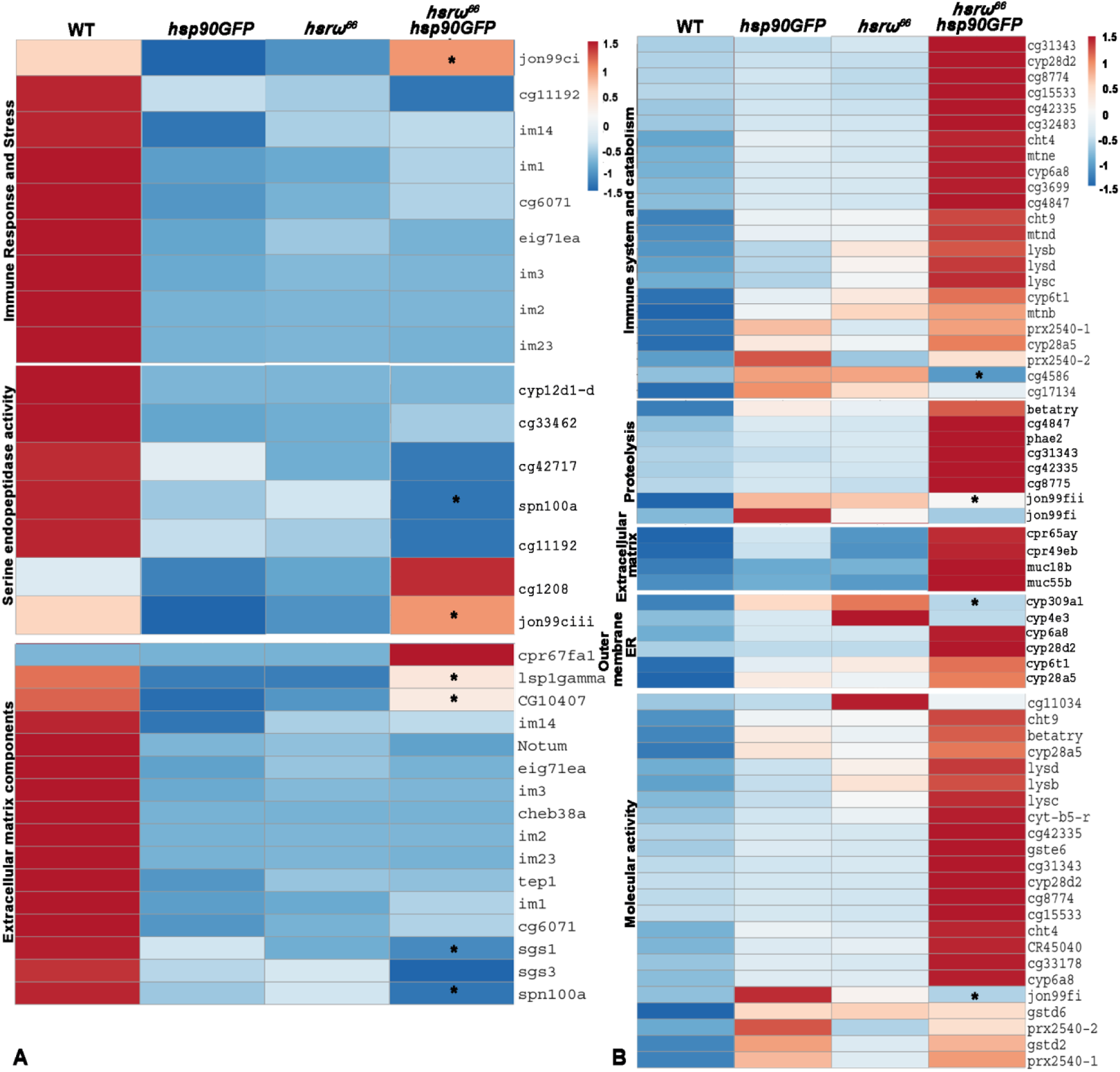
Transcripts of many genes in different pathways are commonly affected in *hsrω*^*66*^ and *Hsp90GFP* larvae with additive effects in *hsrω*^*66*^ *Hsp90GFP* homozygotes. Heat maps, based on FPKM values (Supplementary Table S4, Excel sheets 1 and 2), for different genes (right of each row), grouped according to GO terms (left) in different genotypes (top of each column), which, compared to wild type, are either significantly down- (**A**) or up-regulated (**B**) in both *hsrω*^*66*^ and *hsp90GFP* larvae. Barring a few, marked by asterisks in *hsrω*^*66*^ *Hsp90GFP* column, which do not show significant difference from those in WT, most genes are further down- (**A**) or up- (**B**) regulated, respectively, in *hsrω*^*66*^ *Hsp90GFP*. Colour keys (top middle for **A** and top right for **B)** indicate the arbitrary relative expression on the basis of relative FPKM values.

A gene ontology (GO) term analysis of the genes commonly affected by loss of *hsr*□ transcripts (*hsr*□^*66*^) or over expression of Hsp83 (*Hsp90GFP*) revealed that the down-regulated genes were related mostly to immunity, serine endopeptidase activity and extracellular matrix (Supplementary Table S1 and Fig. 5A). The commonly up-regulated genes in *hsr*□^*66*^ and *Hsp90GFP* larvae were found to be associated with immunity, proteolysis, catabolism, extracellular matrix and outer membrane of endoplasmic reticulum (Supplementary Table S2 and Fig. 5B). Their molecular functions ranged from diverse peptidase (aminopeptidase, metallopeptidase, serine endopeptidase) to hydrolase, oxido-reductase, peroxidase or of antioxidant activities (Supplementary Tables S1 and S2).

Examination of GO terms of the genes commonly affected in *hsr*□^*66*^ and *Hsp90GFP* genotypes (Fig. 5) did not indicate that these genes could be responsible for the unusual phenotypes displayed by *hsr*□^*66*^ *Hsp90GFP* larvae. Moreover, compared to the relatively small number of genes that were commonly affected in *hsr*□^*66*^ and *Hsp90GFP*, the proportion of genes showing unique down- or up-regulation in *hsr*□^*66*^ *Hsp90GFP* larvae, compared to wild type or *hsr*□^*66*^ or *Hsp90GFP* larvae (Fig. 4) was much larger. Thus, ~90% and ~78% of differentially expressed genes were uniquely down- and up-regulated, respectively, in *hsr*□^*66*^ *Hsp90GFP* larvae (Fig. 4A, B). This shows that while substantial reduction of *hsrω* transcripts or over-expression of Hsp83 affected several genes similarly, the effects become dramatically different and more pervasive when the *hsrω* transcript levels are substantially reduced in presence of elevated levels of Hsp83 so that nearly 1850 genes get uniquely affected in *hsr*□^*66*^ *Hsp90GFP* larvae (Fig. 4, Supplementary Table S3).

GO term search for genes uniquely down- or up-regulated in *hsrω*^*66*^ *Hsp90GFP* larvae, when compared to wild type, revealed that a diverse array of important biological pathways were affected in the double mutant genotype (Supplementary Table S3). While the up-regulated genes belonged mostly to metabolic and biosynthesis pathways including energy production and membrane transport, the down-regulated genes included many different pathways (Fig. 6 and Supplementary Table S3). A large number of genes involved in metamorphosis including those involved in ecdysone synthesis (Fig. 6), were significantly down-regulated in *hsrω*^*66*^ *Hsp90GFP.* This may be responsible for the failure of these larvae to pupate. Negative regulators of neuronal development and those responsible for cell morphology during neuronal differentiation were also highly down-regulated in *hsrω*^*66*^ *Hsp90GFP* larvae (Fig. 6), which may contribute to their brain phenotypes. Since Notch signaling is known to be initiator of neuronal differentiation (Baonza and Freeman 2001; Grandbarbe *et al.* 2003; Hämmerle and Tejedor 2007), down-regulation of many of its interactors also seems to contribute to the CNS phenotype. Likewise, down-regulation of genes for salivary gland morphogenesis is reflected in the poor development of SG in *hsrω*^*66*^ *Hsp90GFP* larvae. Genes involved in imaginal disc morphogenesis were also highly down-regulated in *hsrω*^*66*^ *Hsp90GFP* larval transcriptome (Fig. 6).

**Fig. 6.**
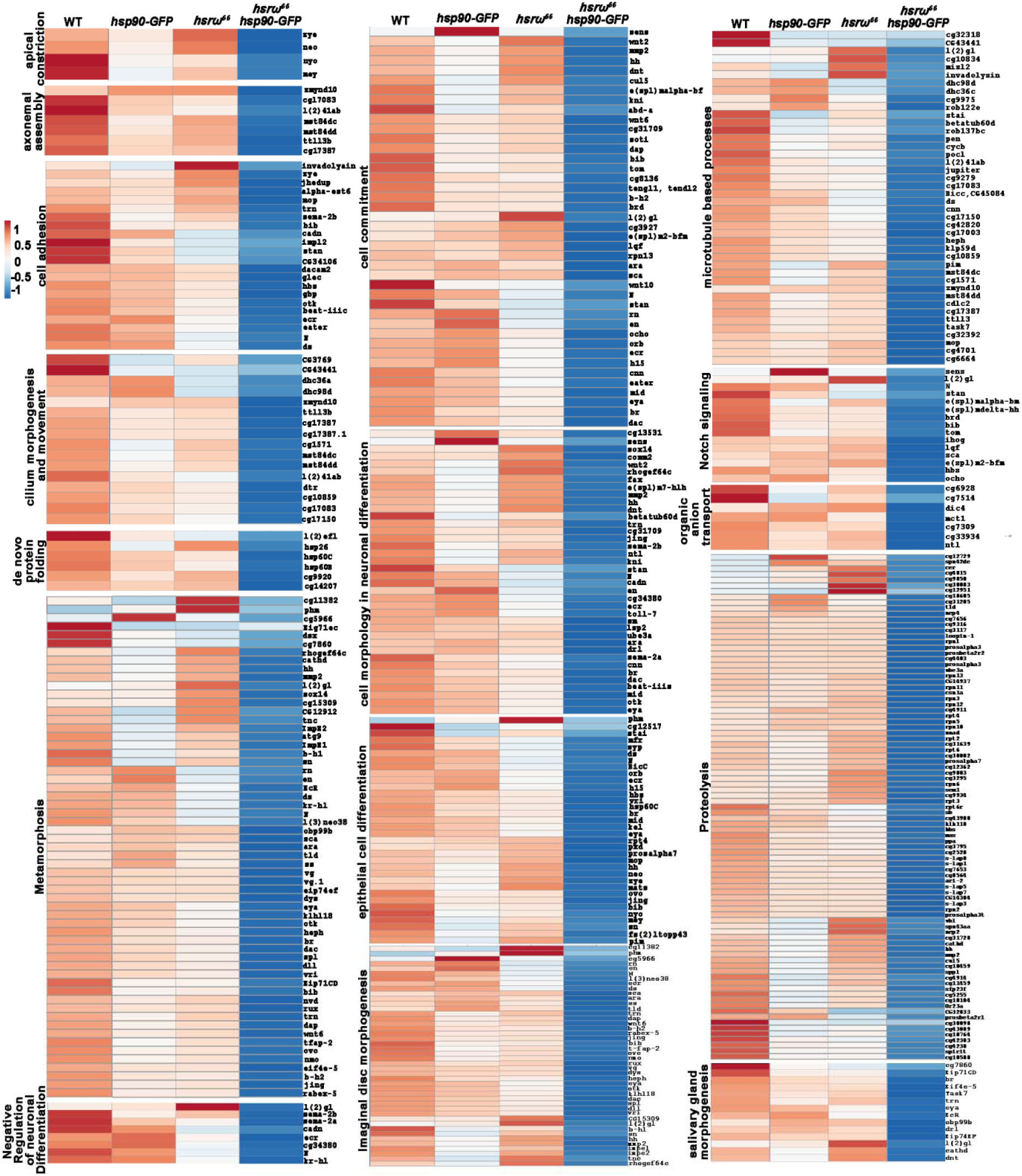
Several genes involved in cell and tissue level organization are exclusively down-regulated in *hsrω*^*66*^ *Hsp90GFP* larvae. Heat maps, based on FPKM values (Supplementary Table S4, Excel sheet 3), of different genes (right of each row) grouped into different GO terms (left) in different genotypes (noted on top of each column) that are significantly down-regulated in *hsrω*^*66*^ *Hsp90GFP*. Colour key (top left) indicates the arbitrary relative expression values on the basis of FPKM values.

Among the genes belonging to cell commitment and morphogenesis pathways, the apico-basal cell polarity determining *l(2)gl* gene (Bilder *et al.* 2000; Calleja *et al.* 2016; Carvalho *et al.* 2015; Hariharan and Bilder 2006; Humbert *et al.* 2008; Humbert *et al.* 2003; Klämbt and Schmidt 1986) was highly down-regulated in *hsrω*^*66*^ *Hsp90GFP* larvae but not in *hsrω*^*66*^ or *Hsp90GFP* larvae (Fig. 7A, C). This is significant in view of the great similarities in many phenotypes seen in the *hsrω*^*66*^ *Hsp90GFP* larvae and those in the *l(2)gl* loss of function mutants. The *l(2)gl* loss of function mutation is well known to result in malignant transformation of brain and imaginal disc cells, under-development of larval SG and PG and a prolonged larval life without pupation (Betschinger *et al.* 2003; Farkas and Mechler 2000; Gateff and Mechler 1989; Ohshiro *et al.* 2000; Roy and Lakhotia 1991). Unlike the *l(2)*gl, levels of *dlg* and *scribble*, two other apico-basal polarity genes, were not significantly affected in any of the genotypes (Fig 7A, C).

**Fig. 7.**
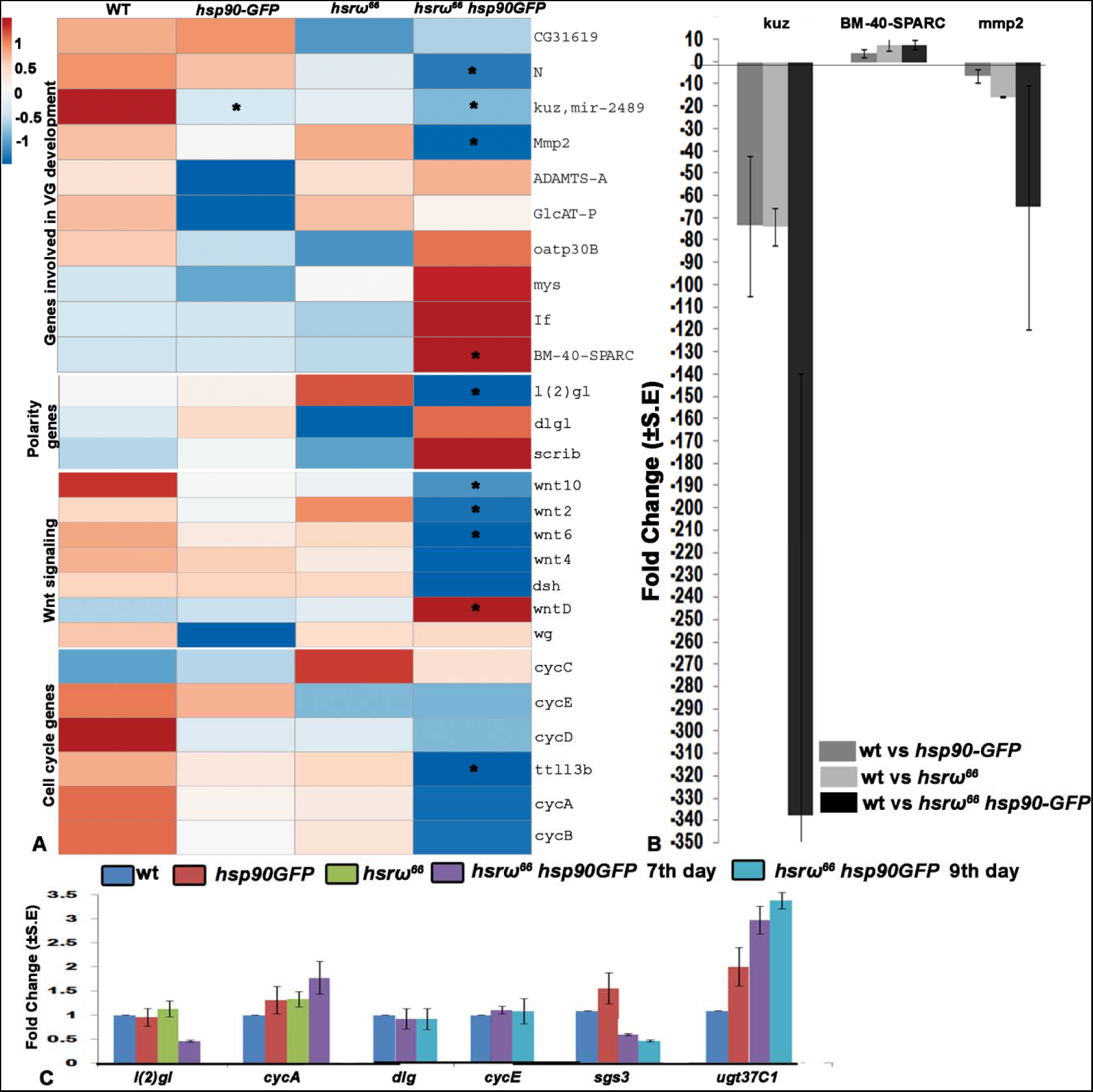
The *hsrω* ^*66*^ *Hsp90GFP* larvae show decrease in level of tumor suppressor gene *l(2)gl* and differential expression of VG shape determining genes. **A** Heat maps, based on FPKM values (Supplementary Table S4, Excel sheet 4), of genes (noted on right of each row) belonging to GO terms relating to development of the VG and cell proliferation (noted on left) in different genotypes (mentioned at top of each column). Genes significantly down-or up-regulated, compared to WT (column 1) are marked by asterisk. **B-C** Histograms showing relative fold changes in levels of transcripts of genes (x-axis) in whole larvae of genotypes (colour keys on top) based on qRT-PCR (**B**) or semi-quantitative RT-PCR (**C**) results.

Changes in levels of many proteolytic pathway genes in *hsrω*^*66*^ *Hsp90GFP* (Fig. 6) may also contribute to the highly elongated VG seen in case of *hsrω*^*66*^ *Hsp90GFP* since an optimal balance of proteolytic activity is required to maintain the structural integrity of neural lamella, the extracellular matrix in neural tissues, which interacts with the underlying glia to shape the *Drosophila* nervous system (Hoang and Chiba 1998; Martinek *et al.* 2008; Meyer *et al.* 2014; Xie and Auld 2011). Examination of genes known to be involved in VG development, revealed that transcripts of genes like *kuz* (Kuzbanian protease) and *mmp2* (an extra-cellular matrix metalloprotease) were significantly down-regulated while *SPARC* (an extra-cellular matrix protein) was up-regulated only in *hsrω*^*66*^ *Hsp90GFP* larvae (Fig. 7A). Interaction between the neural lamella (extra-cellular lamella) and glia shapes the VG and either down-or up-regulation of its components results in elongated nerve cord phenotype (Kato *et al.* 2011; Martinek *et al.* 2008; Meyer *et al.* 2014; Pandey *et al.* 2011; Schmidt *et al.* 2012; Xie and Auld 2011). Expressions of these genes were affected in *Hsp90GFP* as well as *hsrω*^*66*^ but to a much less extent than in *hsrω*^*66*^ *Hsp90GFP* larvae (Fig. 7A, B).

Several genes involved in axonemal assembly, cilium morphogenesis, protein folding, and proteolysis were also significantly down-regulated only in *hsrω*^*66*^ *Hsp90GFP* larvae. Axonemal proteins, the main components of cilia and flagella, form components of primary cilia in eukaryotic cells (Patel-King and King 2016; Satir *et al.* 2010). In higher vertebrates their dysfunction results in a range of ciliopathy syndromes, marked by tumor formation and impairment in neurogenesis (Berbari *et al.* 2009; Guemez-Gamboa *et al.* 2014; Louvi and Grove 2011). Although the chondotonal sensory neurons and sperm are the only tissues that have been reported to have primary cilia in *Drosophila* (Jana *et al.* 2016; Karak *et al.* 2015; Riparbelli *et al.* 2012), recent reports have also implicated roles of the ciliary proteins in development since mutations in some of these gene cause embryonic lethality (Moore *et al.* 2013; Rogowski *et al.* 2009). The ciliary component gene like *ttl3b*, whose mutation results in embryonic lethality (Rogowski *et al.* 2009), and *zmynd10*, mutation in which leads to primary ciliary dyskinesia in both flies and humans (Moore *et al.* 2013), were significantly down-regulated in *hsrω*^*66*^ *Hsp90GFP* larvae (Fig. 6 and Table S4).

The Wnt family regulates neuronal differentiation and cell fate commitment (Alexandre *et al.* 2014; Chiang *et al.* 2009; Inestrosa and Varela-Nallar 2014; Oliva *et al.* 2013). Some members of this family were significantly down–regulated in *hsrω*^*66*^ *Hsp90GFP* larvae (Fig. 7A). Expression of the canonical Wnt signaling pathway genes like *wingless*, a segment polarity gene, and *dsh*, ligand for Wnt signaling (Cohen *et al.* 2008; Reichsman *et al.* 1996; Sato 2006), was not affected in any of the genotypes examined. Similarly, expression of the non-canonical Wnt signaling pathway and planar cell polarity gene *wnt4* (Lim *et al.* 2005; Sato 2006) also did not show any significant change in any of the genotypes. However, several other members of this pathway, like *wnt2, wnt6, wnt10* and *wntD*, showed differential expression in *hsrω*^*66*^ *Hsp90GFP* (Fig. 7A). It is notable that protein products of the down-regulated *wnt2, wnt6, wnt10* genes have serine residues (Herr *et al.* 2012). On the other hand, the *wntD* gene, whose product does not have any serine residue (Ching *et al.* 2008; Herr *et al.* 2012) was up-regulated. Such differential effects may be related to several serine peptidases being affected (Fig. 5) to a much greater extent in *hsrω*^*66*^ *Hsp90GFP* homozygotes.

As a general validation of the RNA seq data, we used qRT-PCR (Fig. 7B) or semi-quantitative RT-PCR (Fig. 7C) to assess levels of transcripts of several genes that were, on the basis of RNA seq data, either not affected (*dlg, CycE* and *CycA*) or were down-regulated (*sgs3, l(2)gl, kuz* and *mmp2*) or up-regulated (*ugt37Ct* and *SPARC*) in *hsrω*^*66*^ *Hsp90GFP* larvae. Results presented in Fig. 7B-C show that there was a complete agreement between the RNA seq and RT-PCR data. This provides a general confidence in the RNA seq data in different genotypes.

### 3.4 Several transcription factor and RNA binding protein encoding genes were uniquely down-regulated in hsrω^66^ Hsp90GFP homozygotes

Since our data revealed wide perturbations in levels of various gene transcripts in *hsrω*^*66*^ *Hsp90GFP* homozygotes, which may result from changes in transcription and/or RNA processing/stability, we examined status of transcription factor (TF) genes in the different genotypes using the list of 996 transcription factors in *Drosophila* (Rhee *et al.* 2014) to identify those whose transcript levels were affected in *hsrω*^*66*^ *Hsp90GFP* homozygotes but not in the other genotypes. It was found that unlike the effect on only a very limited but non-overlapping sets of TF in *hsrω*^*66*^ or *Hsp90GFP* homozygotes, expression of 64 TF genes was significantly (P<0.05) down-or up-regulated in *hsrω*^*66*^ *Hsp90GFP* homozygotes (Fig. 8A and B).

**Fig. 8.**
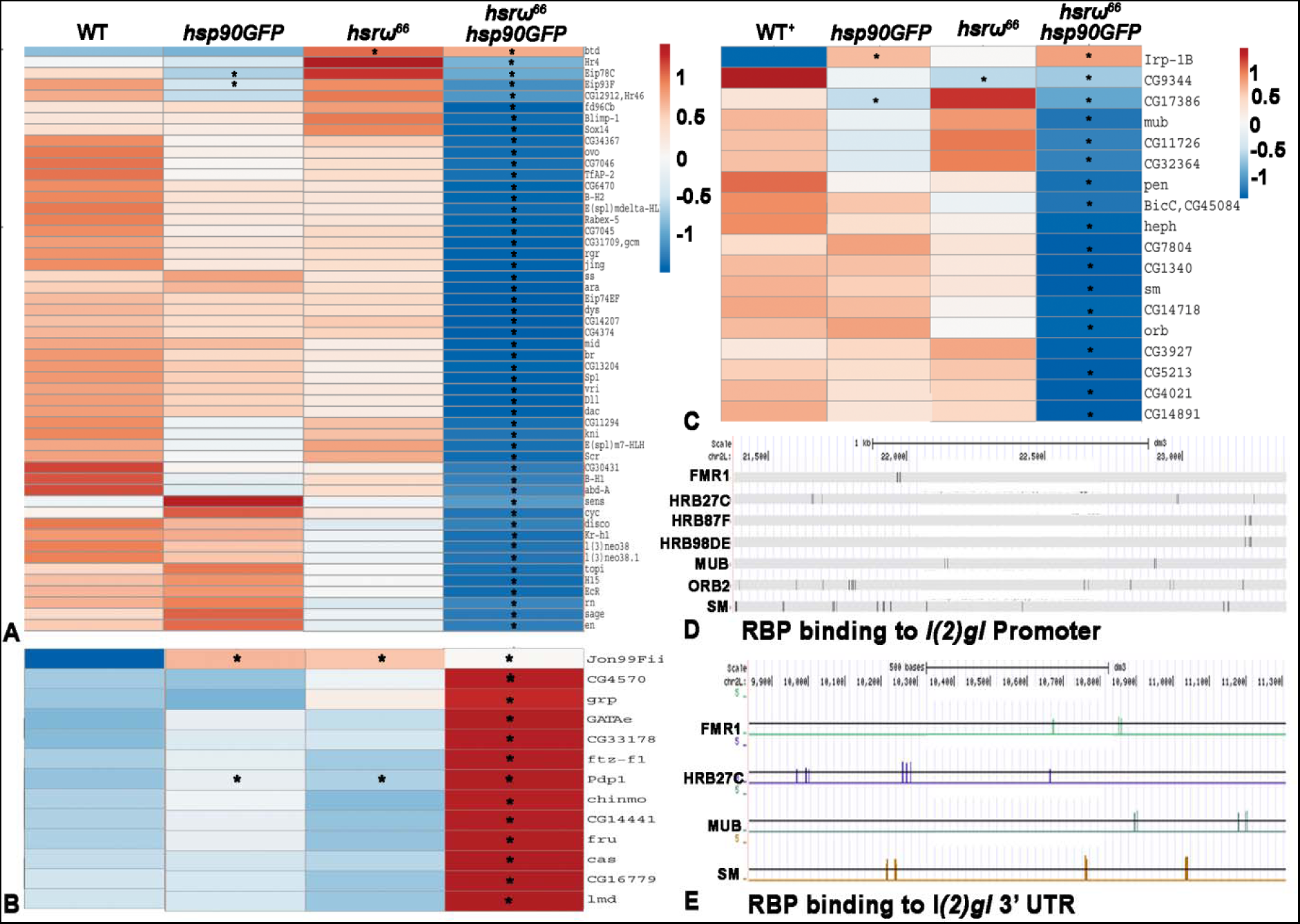
Several transcription factors and RNA binding proteins (RBP) show differential expression in *hsrω*^*66*^ *Hsp90GFP* larvae. **A-B** Heat maps, based on FPKM values (Supplementary Table S4, Excel sheets 5, 6 and 7) of transcription factor genes (noted on right of each row) in different genotypes (noted at top of each column) that were significantly down- (**A**) or up-regulated (**B**) in *hsrω*^*66*^ *hsp83GFP*. **C** Heat maps of FPKM values of RNA binding protein genes (noted on right of each tow) in different genotypes (noted on top of each column) significantly affected in *hsrω*^*66*^ *hsp83GFP.* Asterisks denote significant changes compared to corresponding WT (column 1). The colour keys denote arbitrary values given on basis of relative FPKM values for each gene in the four genotypes. **D-E** Diagrammatic representation of *l(2)gl* promoter (**D**) and 3’UTR (**E**) showing predicted binding of RBPs at the *l(2)gl* gene.

We also checked the levels of RNA binding proteins (RBP) listed in RBPDB (http://rbpdb.ccbr.utoronto.ca/) and found that out of the 259 genes, 18 showed differential expression in *hsrω*^*66*^ *Hsp90GFP* homozygotes (Fig. 8C). It was interesting that of these only one showed a similar significant (P<0.05) down-regulation in *hsrω*^*66*^ while in *Hsp90GFP* homozygous larvae two genes showed changes similar to those in *hsrω*^*66*^ *Hsp90GFP* homozygotes (marked by asterisks in Fig. 8C). Surprisingly, levels of transcripts for none of the known omega speckle associated hnRNPs and those whose intracellular distribution is affected in *hsrω*^*66*^ cells (Lakhotia *et al.* 2012; Mallik and Lakhotia 2011; Piccolo *et al.* 2017; Piccolo and Yamaguchi 2017; Singh and Lakhotia 2015) were significantly altered in any of the three mutant genotypes examined. However, *heph* and *sm*, which encode members of the hnRNP-L family, were down-regulated in *hsrω*^*66*^ *Hsp90GFP* homozygotes. Sm is predicted to bind to both *l(2)gl* promoter as well as its 3’UTR (Fig. 8D, E). These two hnRNP family proteins, like the other hnRNPs (Lakhotia 2011; Piccolo *et al.* 2017; Piccolo and Yamaguchi 2017), are expected to be associated with the omega speckles although this has not yet been experimentally demonstrated.

A bioinformatic analysis suggested that some of the affected RBPs have potential binding sites in the *l(2)gl* promoter and/or 3’UTR (Fig. 8D, E). We also found that two important transcription factors dFus (not shown) and dFMRP/FMR1 (Fig 8D) are predicted to bind to *l(2)gl* promoter. This is significant since Fus interacts directly with hsr*ω* RNA (Piccolo *et al.* 2017) while one of the interactors of FMR1, TDP43 also shows physical interaction with hsr*ω* RNA (Piccolo et al 2018). Interestingly, Hrb87F and Hrb98DE, which are two of the principle hnRNPs known to be associated with *hsrω* RNA (Prasanth *et al.* 2000; Singh and Lakhotia 2016), also show potential binding sites with the *l(2)gl* promoter (Fig. 8D).

Analysis of the available chip-seq data for Hsp90 (Sawarkar *et al.* 2012) and that of one of its interactor, Trithorax (Tariq *et al.* 2009), which is involved in chromatin remodeling and thus influences gene expression (Kassis *et al.* 2017; Ringrose 2017; Schuettengruber *et al.* 2017) revealed that both these proteins bind to *l(2)gl, kuz* and *hsrω* genes (Table 5). Their binding sites in the *l(2)gl* gene promoter are overlapping, which suggests that altered levels of Hsp83 can affect transcription at this locus. Additionally, the *hsrω* gene locus also shows strong Hsp83 binding in Chip-seq data as well as thorough immunostaining (Table 5) (Sawarkar *et al.* 2012; Tariq *et al.* 2009).

**Table 5.**
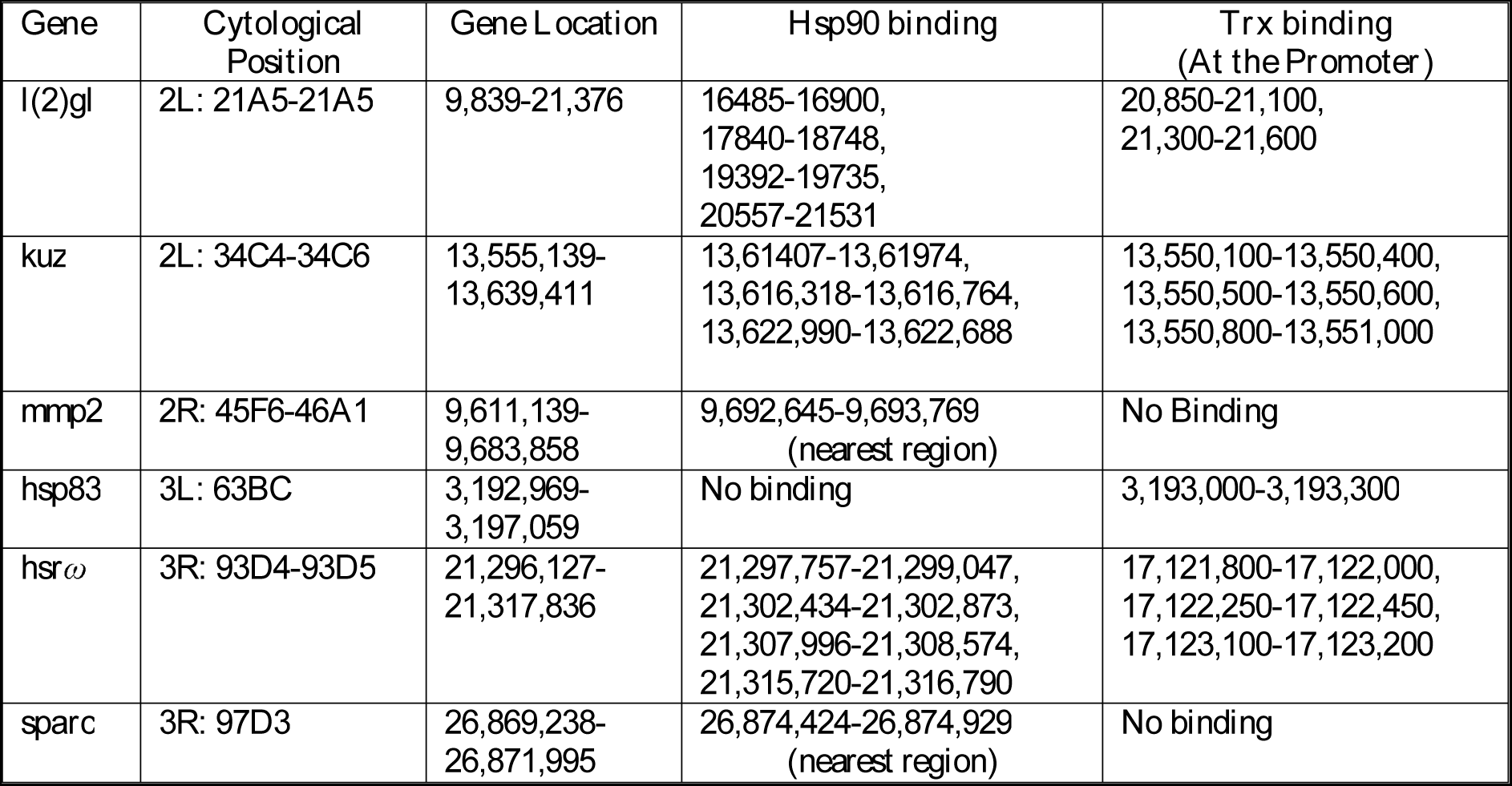
Hsp83 and Trithorax proteins bind with promoters of several genes implicated in this study (based on Chip-seq data of Savarkar et al (2012))

| **Gene** | **Cytological Position** | **Gene Location** | **Hsp90 binding** | **Trx binding**<br>**(At the Promoter)** |
| --- | --- | --- | --- | --- |
| *l(2)gl* | 2L: 21A5-21A5 | 9,839-21,376 | 16485-16900,<br>17840-18748,<br>19392-19735,<br>20557-21531 | 20,850-21,100,<br>21,300-21,600 |
| *kuz* | 2L: 34C4-34C6 | 13,555,139-13,639,411 | 13,61407-13,61974,<br>13,616,318-13,616,764,<br>13,622,990-13,622,688 | 13,550,100-13,550,400,<br>13,550,500-13,550,600,<br>13,550,800-13,551,000 |
| *mmp2* | 2R: 45F6-46A1 | 9,611,139-9,683,858 | 9,692,645-9,693,769<br>(nearest region) | No Binding |
| *hsp83* | 3L: 63BC | 3,192,969-3,197,059 | No binding | 3,193,000-3,193,300 |
| *hsrw* | 3R: 93D4-93D5 | 21,296,127-21,317,836 | 21,297,757-21,299,047,<br>21,302,434-21,302,873,<br>21,307,996-21,308,574,<br>21,315,720-21,316,790 | 17,121,800-17,122,000,<br>17,122,250-17,122,450,<br>17,123,100-17,123,200 |
| *sparc* | 3R: 97D3 | 26,869,238-26,871,995 | 26,874,424-26,874,929<br>(nearest region) | No binding |

### 3.5 The Sp/CyO chromosomes do not seem to contribute to the above noted gene expression changes in Sp/CyO; hsrω^66^ Hsp90GFP larvae

Since in the above comparisons, the *hsrω*^*66*^ *Hsp90GFP* samples differed from the other three genotypes (*Oregon R*^*+*^ or WT, +/+; *hsrω*^*66*^ *Hsp90GFP*) in carrying *Sp/CyO* chromosomes instead of the wild type chromosome 2 and since the *+/+; hsrω*^*66*^ *Hsp90GFP* died as very early larvae while the *Sp/CyO; hsrω*^*66*^ *Hsp90GFP* larvae survived to 3^rd^ instar stage, it remained possible that the *Sp/CyO* chromosomes may also contribute to the unusual phenotypes displayed by the rare surviving *Sp/CyO*; *hsrω*^*66*^ *Hsp90GFP* larvae. Since transcriptomes of *+/+*; *hsrω*^*66*^ *Hsp90GFP* and *Sp/CyO*; *hsrω* ^*66*^ *Hsp90GFP* larvae could not be compared directly because none of the former survived to 3^rd^ instar stage, we compared transcriptomes of *+/+; hsrω*^*66*^ and *Sp/CyO; hsrω*^*66*^ 3^rd^ instar larvae to get some information about effects of *Sp/CyO* chromosomes.

The transcriptome data revealed that expression of only a small number of genes was different between *+/+; hsrω*^*66*^ and *Sp/CyO; hsrω* ^*66*^ larvae. On the basis of significant *p*-value differences (*p* <0.05), 146 and 211 genes were down-regulated in *+/+; hsrω*^*66*^ and *Sp/CyO; hsrω* ^*66*^, respectively, when compared to WT; of these 85 were common between the two sets. Similarly,335 and 407 genes were up-regulated in *+/+; hsrω*^*66*^ and *Sp/CyO; hsrω* ^*66*^, respectively, when compared to WT; of these 214 were common between the two sets. In order to reduce the probability of false-positives, we also compared the numbers of genes showing differential expression on the *q*-value significance. This comparison revealed 70 and 42 genes to be down-regulated, compared to WT, in *+/+; hsrω*^*66*^ and *Sp/CyO; hsrω* ^*66*^, respectively, with 24 being common (Fig. 9A). Similarly, 77 and 97 genes were up-regulated, compared to WT, in *+/+;ω* and *Sp/CyO; hsrω*, respectively with 39 being common (Fig. 9B). A comparison between *+/+; hsrω*^*66*^ and *Sp/CyO; hsrω* ^*66*^ transcriptomes revealed much smaller difference between the two with only 26 genes being up-regulated and 11 down-regulated in *Sp/CyO; hsr ω*^*66*^ compared to *+/+; hsrω*^*66*^. These are listed in Figs. 9C and D. We examined expression levels of these genes in *Sp/CyO; hsrω*^*66*^ *Hsp90GFP* larvae. Interestingly, only three genes (*Cht4, Yp3* and *CG6788*) were similarly down regulated in *Sp/CyO; hsrω*^*66*^ and *Sp/CyO; hsrω* ^*66*^ *Hsp90GFP*. On the other hand, two genes (*Cpr47Eg* and *CG14219*) that were down-regulated in *Sp/CyO; hsrω*^*66*^ larvae, were actually up-regulated in *Sp/CyO; hsr ω*^*66*^ *Hsp90GFP*. None of the26 up-regulated gene in *Sp/CyO; hsrω* ^*66*^ larvae showed significant difference in their expression levels in WT and *Sp/CyO; hsrω*^*66*^ *Hsp90GFP* larvae.

**Fig. 9.**
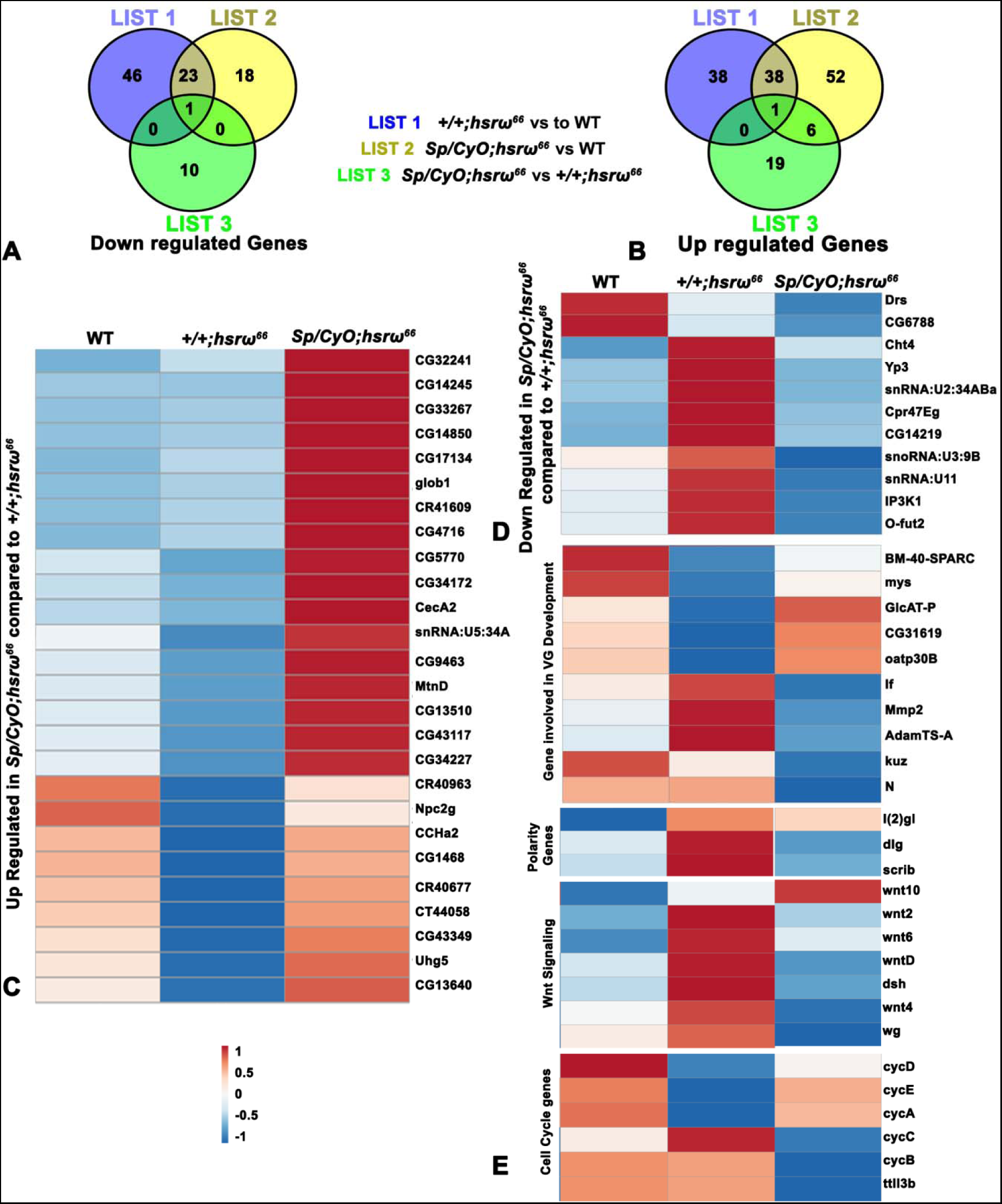
Fig. 9. *+/+; hsrω***^*66*^ and *Sp/CyO; hsr****ω*^*66*^ larval transcriptomes show limited differences. **A-B** Venn diagrams showing the numbers of genes that are significantly down- (**A**) or up-regulated (**B**) in *+/+; hsrω*^*66*^ (List 1) and *Sp/CyO; hsrω*^*66*^ (List 2) compared to wild type (WT) and in *Sp/CyO; hsrω*^*66*^ compared to *+/+; hsrω*^*66*^ (List 3). **C-D** Heat maps, based on FPKM values (Supplementary Table S4, Excel sheets 8 and 9) for genes (noted on right) that are up- (**C**) or down-regulated (**D**) in *Sp/CyO; hsrω*^*66*^ compared to *+/+; hsrω*^*66*^ in different genotypes (noted on top of each column). **E** Heat maps of genes (noted on right), based on FPKM values (Supplementary Table S4, Excel sheet 10), belonging to different GO terms (noted on left) involved in development of the VG and cell proliferation in different genotypes (noted at top of each column). The colour key (below **C**) applies to **C**-**F** and denotes arbitrary values given on basis of relative FPKM values for each gene in the three genotypes.

None of the different pathway genes shown in Fig. 7A, whose altered expression in *Sp/CyO; hsrω*^*66*^ *Hsp90GFP* appeared to be critical for their phenotypes, exhibited differential expression (neither on *p* nor on *q* values) between *+/+; hsrω*^*66*^ and *Sp/CyO; hsrω*^*66*^ (Fig 9E). It is significant to note that l*(2)gl, kuz* and *SPARC* genes did not show any significant difference in their expression between *+/+; hsrω*^*66*^ and *Sp/CyO; hsrω*^*66*^. Likewise, the various RBP and TF genes, which were affected in *Sp/CyO; hsrω*^*66*^ *Hsp90GFP* larvae, did not show any difference in theirexpression between *+/+; hsrω*^*66*^ and *Sp/CyO; hsrω*^*66*^ larvae.

Together, therefore, the *Sp/CyO* chromosomes do not appear to contribute to the unusual phenotypes exhibited by *Sp/CyO; hsrω*^*66*^ *Hsp90GFP* larvae, except that these chromosomes somehow delay the death of larvae with elevated Hsp83 in a background of grossly depleted *hsrω* transcripts after the organisms reach third instar stage.

## 4. Discussion

We undertook this study to further examine the functional interaction between ubiquitously expressed and stress-inducible *hsrω* lncRNA and *Hsp83* genes in view of the several earlier results (Lakhotia and Ray 1996; Morcillo *et al.* 1993; Tariq *et al.* 2009) that suggested possible interaction/association between their products. A near absence of *hsrω* transcripts in *hsrω*^*66*^ homozygotes causes some degree of embryonic and late-pupal stage lethality but, as seen in a previous study (Lakhotia *et al.* 2012), does not affect pupation of the larvae that hatch; present study), although, as noted here, they show a slight delay in larval development. Likewise, the Hsp83 over-expressing transgenic line *Hsp90GFP* does not show any obvious developmental defects (Tariq *et al.* 2009), except a small delay in larval development noted in this study. Therefore, the very high embryonic death and prolonged larval life followed by death without pupation of the rare surviving *hsrω*^*66*^ *Hsp90GFP* homozygous larvae was unexpected, more so because the *hsrω*^*66*^ *Hsp90GFP/TM6B* double heterozygotes did not show any apparent developmental defects or other phenotypic changes.

The common down- or up-regulation of a significant number of genes in *hsrω*^*66*^ or *Hsp90GFP* homozygous larvae, and the additive/synergistic effect seen in expression of many of these genes in *hsrω*^*66*^ *Hsp90GFP* homozygotes, revealed by our whole larval transcriptome analysis, indicate that products of these two genes indeed participate in common pathways and therefore, mutant alleles of the two may be expected to affect the phenotype when together. Interacting genes generally show some phenotypic consequence in heterozygous combination as well. Intriguingly however, the *hsrω*^*66*^ *Hsp90GFP/TM6B* heterozygotes did not provide any indication about the severe consequences when the *hsrω*^*66*^ *Hsp90GFP* combination becomes homozygous. The *hsp83* transcripts in these heterozygotes were only marginally less than in *Hsp90GFP* homozygotes (Fig. 1). The *hsrω* transcript levels in *hsrω*^*66*^ heterozygotes were reduced, but this level of reduction apparently did not significantly affect the survival of the *hsrω*^*66*^ *Hsp90GFP* heterozygotes. This is further confirmed by our finding that ubiquitous expression of *hsrωRNAi* transgene, which also reduced levels of different *hsrω* transcripts but did not cause any enhanced lethality when expressed in *Hsp90GFP* homozygotes. Thus, only when the levels of *hsrω* transcripts are reduced below certain minimal levels in a background of elevated Hsp83 levels, the severe adverse phenotypic effects manifest themselves.

Since the unusual phenotypes were seen only in *Sp/CyO; hsrω*^*66*^ *Hsp90GFP* but not in *+/+; hsrω*^*66*^ *Hsp90GFP* genotype, it remained possible that the *Sp/CyO* chromosomes may also contribute to the unusual late larval phenotypes. However, the *Sp/CyO* chromosome combination actually seems to reduce the severity of phenotype since while *+/+; hsrω*^*66*^ *Hsp90GFP* homozygous larvae completely failed to survive beyond the 1st or 2nd instar stage, at least some of the *Sp/CyO; hsrω*^*66*^ *Hsp90GFP* larvae reached the third instar stage before dying. Our comparison of transcriptomes of *+/+; hsrω*^*66*^ and *Sp/CyO; hsrω*^*66*^ showed only a limited number of genes to be affected by the presence of *Sp/CyO* chromosomes in *hsrω*^*66*^ background. Since the set of genes differentially expressed between *+/+; hsrω*^*66*^ and *Sp/CyO; hsrω*^*66*^ did not include any of those that were uniquely affected in *Sp/CyO; hsrω*^*66*^ *Hsp90GFP* larvae, the combined effect of homozygosity for *hsrω*^*66*^ *Hsp90GFP* alleles seems to be prime cause for the observed larval phenotypes.

The larval phenotypes, viz., prolonged larval life, bulbous and transparent appearance of aged larvae, disorganized and clumped imaginal discs, small size of endoreplicating tissues like salivary glands, prothoracic glands, gut, Malpighian tubules etc, and the enlarged brain lobes, displayed by *hsrω*^*66*^ *Hsp90GFP* homozygotes are striking phenocopy of those seen in larvae homozygous for the *l(2)gl* loss of function alleles (Farkas and Mechler 2000; Gateff and Mechler 1989; Merz *et al.* 1990; Roy and Lakhotia 1991). The severely reduced levels of *l(2)gl* transcripts, shown by our RNA seq and RT-PCR data, in *hsrω*^*66*^ *Hsp90GFP* homozygotes, but not in *hsrω*^*66*^ *Hsp90GFP/TM6B* heterozygotes or in *hsrω*^*66*^ (+/+; *hsrω*^*66*^ or *Sp/CyO; hsrω*^*66*^) or *Hsp90GFP* homozygotes, suggests that the unusual phenotypes of *hsrω*^*66*^ *Hsp90GFP* homozygous larvae are indeed a consequence of the substantially down-regulated expression of the apico-basal polarity *l(2)gl* gene. The elongated VG phenotype displayed by the surviving *hsrω*^*66*^ *Hsp90GFP* 3rd instar larvae is not a characteristic feature of the *l(2)gl* mutant larvae. This seems to result from the altered transcript levels of genes like *kuz, mmp2, SPARC* etc that are known to regulate shaping of the embryonic and larval VG (Meyer *et al.* 2014). These genes were also differentially expressed only in *hsrω*^*66*^ *Hsp90GFP* homozygotes, but not in *Sp/CyO; hsrω*^*66*^ *Hsp90GFP/TM6B* heterozygotes or in *hsrω*^*66*^ or *Hsp90GFP* single homozygotes. Since neither Hsp83 nor the *hsrω* lncRNAs directly function as transcription factors or as regulators of RNA turnover, the observed alterations in transcripts of *l(2)gl, kuz, mmp2, SPARC* and the large number of other genes in *hsrω*^*66*^ *Hsp90GFP* homozygotes seem to be indirect consequences of their altered levels.

The ubiquitously expressed Hsp83, besides its more commonly known role as a protein folding chaperone (Arya *et al.* 2007; Li *et al.* 2007; Rutherford and Lindquist 1998), also regulates i) several signaling pathways through binding with other proteins (Cutforth and Rubin 1994; DeFranco and Csermely 2000; Segnitz and Gehring 1997), ii) organization of cytoskeleton (Fostinis et al 1992, Taiyab and Rao 2011), iii) centrosome function (de Cárcer *et al.* 2001; Lange *et al.* 2000), iv) activities of a large number of genes through its roles in biogenesis of piwiRNA, miRNA, sno/sn/telomerase RNAs etc (Boulon *et al.* 2010; Dittmar and Sen 2018; Gangaraju *et al.* 2011; Iwasaki *et al.* 2015; Miyoshi *et al.* 2010; Olivieri *et al.* 2012; Specchia *et al.* 2010; Zhao *et al.* 2008), and v) local chromatin organization through direct binding at many gene sites (Mazaira *et al.* 2016; Sawarkar *et al.* 2012; Tariq *et al.* 2009). Binding of Hsp83 at many of these chromosome sites overlaps with that of Trx, an important regulator of developmental gene expression through chromatin remodeling (Mazaira *et al.* 2016; Ringrose 2017; Sangster *et al.* 2003; Sawarkar *et al.* 2012; Tariq *et al.* 2009). Interestingly, Hsp83 as well as Trx proteins bind at promoters of genes like *hsp83, l(2)gl* and *hsrω* (Supplementary Table 4). Thus elevated levels of Hsp83 can potentially affect the *l(2)gl* transcription. However, since *l(2)gl* transcripts, like those of many others, were severely down-regulated only when Hsp83 was over-expressed in the background of near absence of *hsrω* transcripts, additional factors, are affected by *hsrω* transcript levels, are needed for the synthetic action of elevated levels of Hsp83 and depleted *hsrω* transcripts on *l(2)gl* and the other affected genes.

The globally expressed *hsrω* gene produces multiple nuclear and cytoplasmic non-coding transcripts (Lakhotia 2011, 2016; Lakhotia 2017). The larger nuclear *hsrω* transcripts are essential for the assembly of nucleoplasmic omega speckles harbouring a variety of hnRNPs and some other RNA binding proteins so that in the absence or reduced or elevated levels of the *hsrω* nuclear transcripts, the associated hnRNPs and other proteins lose their characteristic speckled nucleoplasmic distribution, which affects their sub-cellular localization, dynamic mobility and availability (Lakhotia 2017; Mallik and Lakhotia 2011; Piccolo *et al.* 2017; Piccolo and Yamaguchi 2017; Prasanth *et al.* 2000; Singh and Lakhotia 2015).

Functions of various hnRNPs range from transcriptional regulation to post-transcriptional processing, transport, spatial localization and degradation of different RNAs in a tissue and transcript-specific manner (Appocher *et al.* 2017; Chaudhury *et al.* 2010; Coyne *et al.* 2015; Estes *et al.* 2008; Han *et al.* 2010; Piccolo *et al.* 2014; Romano *et al.* 2016; Specchia *et al.* 2017; Zhang *et al.* 2017). Since the levels of transcripts encoding most of the hnRNPs and other omega speckle associated RBPs were not found to be altered in *hsrω*^*66*^ or *Hsp90GFP* or *hsrω*^*66*^ *Hsp90GFP* homozygotes, the observed effects on expression of various genes in genotypes with reduced levels of *hsrω* transcripts are likely to be consequences of the altered dynamics and spatial distribution of the omega speckle associated hnRNPs and other proteins. Activity at the *hsrω* locus affects poly-ADP-ribosylation of hnRNPs (Ji and Tulin 2009, 2013), which in turn can affect many downstream events like splicing and turnover of RNA. Down-regulation of *hsrω* transcripts also causes hnRNPs like dFUS (Caz) to move to cytoplasm where it shows greater interaction with dFMRP (Piccolo and Yamaguchi 2017). The TDPH/TDP-43 also physically associates with hsr*ω* nuclear RNAs (Piccolo *et al.* 2018) and alterations in poly-ADP-ribose binding affects its liquid-phase separation properties and sub-cellular localization (McGurk *et al.* 2018). The TBPH/TDP-43 interacts with FMRP and together they affect mRNA metabolism at levels of transcription, pre-mRNA splicing, mRNA stability, miRNA biogenesis, transport and translation (Coyne *et al.* 2015; Estes *et al.* 2008; Romano *et al.* 2016; Specchia *et al.* 2017). Thus, altered localizations of dFUS and TDP-43 following depletion of *hsrω* lncRNAs would affect their interaction with dFMRP and other hnRNPs (Romano *et al.* 2016), resulting in variable consequences for different transcripts. The *hsrω* transcripts also interact with chromatin remodeling components like ISWI, CBP300 etc and with nuclear matrix components like Megator, SAFB, Lamin etc (Lakhotia 2017; Onorati *et al.* 2011; Piccolo and Yamaguchi 2017). In addition, the *hsrω* gene also shows genetic interaction with Ras- and JNK-signaling and with caspase-dependent apoptotic pathways (Lakhotia 2017; Ray and Lakhotia 2017). In agreement with such wide-ranging interactions of the *hsrω* lncRNAs, our transcriptomic analysis in *hsrω* ^*66*^ homozygous larvae indeed revealed changes in a large number of genes associated with diverse cellular activities.

Our bioinformatic finding that dFUS and dFMRP may bind with the *l(2)gl* promoter (Fig. 8D) is significant. Moreover, dFMRP is also reported to bind with *l(2)gl* transcripts (Georgieva *et al.* 2012). Therefore, it is likely that altered localization and activities of proteins like dFUS, dFMRP, TDP-43 and other hnRNPs can affect levels of transcripts of genes like *l(2)gl* etc through reduced transcription and/or reduced stability of the transcripts. However, since neither the near absence of *hsrω* transcripts in *hsrω* ^*66*^ nor the enhanced levels of Hsp83 in *Hsp90GFP* homozygotes individually affected transcript levels of genes like *l(2)gl, kuz, mmp2, SPARC* etc, which we believe to be majorly responsible for the unusual phenotypes of the surviving *hsr ω*^*66*^ *Hsp90GFP* homozygotes, the synergistic effects of simultaneous perturbations in *hsrω* lncRNA and Hsp83 levels on cellular regulatory networks seem to be critical for activities of these and many other genes. It is significant in this context that Hsp90 and diverse hnRNPs, including FUS and TDP-43, co-localize and/or directly interact in cells and modulates diverse activities at transcriptional and/or post-transcriptional levels (Chi *et al.* 2018; Ford *et al.* 2002; Jinwal *et al.* 2012; Lackie *et al.* 2017; Zhang *et al.* 2006). Obviously, elevation in levels of Hsp83 in a background where the various hnRNPs are mis-localized, would severely and varyingly impinge upon the diverse interactions between Hsp83 and hnRNPs with unexpected consequences for the gene expression profiles, as indeed seen in our study.

In view of the above considerations, we speculate that the action of Hsp83 and/or Trx on transcription through binding on promoter of genes like *l(2)gl* involves specific interaction with some of the *hsrω* transcript associated RBPs, and vice-versa, so that their misbalanced interactions in double mutants can block efficient transcription of the gene or stability of the transcripts. Potential binding of Hrb87F, and Hrb98DE and other omega speckle associated proteins on the *l(2)gl* promoter and/or 3’UTR of its mRNA may play significant roles in such interactions. Furthermore, the Lgl protein is reported to interact with dFMRP and mRNAs, including *l(2)gl* mRNA, and with other ribonucleoprotein complexes (Zarnescu *et al.* 2005). It is possible that besides the diverse roles of dFMRP, TBPH/TDP-43 and Lgl in regulating activities of a large number of other genes, Lgl may also have an auto-regulatory role so that its absence, in conjunction with mis-localized dFMRP and TBPH/TDP-43 etc can further inhibit *l(2)gl* transcription and or can affect stability of the *l(2)gl* transcripts. Hsp83 associates with *hsrω* transcripts in nucleoplasm and as suggested above, this may facilitate its association with other RNPs and thus with gene promoters. Excess of Hsp83, severely depleted levels of *hsrω*transcripts and mis-localized proteins like dFUS, dFMRP, TDP-43 etc, as in *hsrω*^*66*^ *Hsp90GFP* homozygotes, together can inhibit or up-regulate transcription and/or instability of transcripts of a large number of genes, depending upon repressive or stimulatory effect of the Hsp83 and associated factors on a given gene’s expression. Such widespread perturbations would lead to early death as seen in *+/+; hsrω*^*66*^ *Hsp90GFP* homozygotes. Severe down-regulation of genes like *l(2)gl, kuz, mmp2* etc would lead to the phenotypes seen in the rare surviving *Sp/CyO; hsrω*^*66*^ *Hsp90GFP* late larvae. Further studies are needed to decipher the specific underlying mechanisms.

The present study, besides confirming interactions between *hsrω* transcripts and Hsp83 during development, highlights that perturbation in interactions between ubiquitously expressed coding and non-coding genes can result in dramatically severe consequences, including initiation of timorous growth by down-regulation of important tumor suppressor gene like *l(2)gl*.

## Acknowledgements

We thank the Bloomington *Drosophila* Stock Ctr and Dr. Stephen W. Mckechnie (Australia) and Dr Renato Paro (Switzerland) for providing fly stocks. We thank Developmental Studies Hybridoma Bank (DSHB, Iowa, USA) for anti-Wingless and anti-Dlg, and Prof. Robert Tanguay (Canada) for anti-Hsp83 antibodies. We thank the Department of Biotechnology, Govt. of India (New Delhi) and the Indian Council of Medical Research (New Delhi) for supporting this research. We also thank the Centre of Advanced Studies in Department of Zoology, DBT-BHU Interdisciplinary School of Life Sciences and the Centre of Genetic Disorders (CGD) at BHU for various facilities. Special thanks to Dr Amit Chaurasia of Premas Biotech, CGD, for RNA-sequencing. We acknowledge the Department of Science & Technology, Govt. of India (New Delhi) and the Banaras Hindu University for Confocal Microscopy facility. We thank Dr. Yashvant Patel in our lab for help in RNA sequence analysis.

## Author contributions

MR and SCL planned experiments, analyzed results and wrote the manuscript. MR carried out most of the experimental work and data collection while SA and SS helped in some of the experimental work relating to larval phenotypes and semi-quantitative RT-PCR.

## Data availability

The NGS data for RNA-sequencing in the first set of genotypes (*Oregon R*^*+*^ (WT), *Sp/CyO; hsrω*^*66*^, *Sp/CyO; Hsp90GFP, Sp/CyO; hsrω*^*66*^ *Hsp90GF*) are deposited at GEO (http://www.ncbi.nlm.nih.gov/geo/) with accession no. GSE116476. RNA-seq data, while accession no. for the second set of genotypes (WT and *+/+; hsrω*^*66*^) is GSE120077.

## References

Aggarwal SK, King RC 1969 A comparative study of the ring glands from wild type and 1 (2) gl mutant Drosophila melanogaster. J. Morphology 129 171-199

Alexandre C, Baena-Lopez A, Vincent J-P 2014 Patterning and growth control by membrane-tethered Wingless. Nature 505 180

Appocher C, Mohagheghi F, Cappelli S, Stuani C, Romano M, et al. 2017 Major hnRNP proteins act as general TDP-43 functional modifiers both in Drosophila and human neuronal cells. Nucleic Acids Research 45 8026–8045

Arya R, Mallik M, Lakhotia S 2007 Heat shock genes - integrating cell survival and death. J. Biosciences 32 595–610

Bandura JL, Jiang H, Nickerson DW, Edgar BA 2013 The molecular chaperone Hsp90 is required for cell cycle exit in Drosophila melanogaster. PLoS Genetics 9 e1003835

Baonza A, Freeman M 2001 Notch signalling and the initiation of neural development in the Drosophila eye. Development 128 3889–3898

Basto R, Gergely F, Draviam VM, Ohkura H, Liley K, et al. 2007 Hsp90 is required to localise cyclin B and Msps/ch-TOG to the mitotic spindle in Drosophila and humans. J. Cell Science 120 1278–1287

Bello BC, Izergina N, Caussinus E, Reichert H 2008 Amplification of neural stem cell proliferation by intermediate progenitor cells in Drosophila brain development. Neural Development 3 5

Berbari NF, O’Connor AK, Haycraft CJ, Yoder BK 2009 The primary cilium as a complex signaling center. Current Biology 19 R526–R535

Berger C, Kannan R, Myneni S, Renner S, Shashidhara L, et al. 2010 Cell cycle independent role of Cyclin E during neural cell fate specification in Drosophila is mediated by its regulation of Prospero function. Developmental Biology 337 415–424

Betschinger J, Mechtler K, Knoblich JA 2003 The PAR complex directs asymmetric cell division by phosphorylating the cytoskeletal protein Lgl. Nature 422 326

Bilder D, Li M, Perrimon N 2000 Cooperative regulation of cell polarity and growth by Drosophila tumor suppressors. Science 289 113–116

Boulon S, Pradet-Balade B, Verheggen C, Molle D, Boireau S, et al. 2010 HSP90 and its R2TP/Prefoldin-like cochaperone are involved in the cytoplasmic assembly of RNA polymerase II. Molecular Cell 39 912–924

Calleja M, Morata G, Casanova J 2016 Tumorigenic properties of Drosophila epithelial cells mutant for lethal giant larvae. Developmental Dynamics 245 834–843

Carbajal M, Valet J, Charest P, Tanguay R 1990 Purification of Drosophila Hsp 83 and immunoelectron microscopic localization. European Joutnal Cell Biology 52 147–156

Carvalho CA, Moreira S, Ventura G, Sunkel CE, Morais-de-Sá E 2015 Aurora A triggers Lgl cortical release during symmetric division to control planar spindle orientation. Current Biology 25 53–60

Chaudhury A, Chander P, Howe PH 2010 Heterogeneous nuclear ribonucleoproteins (hnRNPs) in cellular processes: Focus on hnRNP E1’s multifunctional regulatory roles. Rna 16 1449–1462

Chen Y, Schnetz MP, Irarrazabal CE, Shen R-F, Williams CK, et al. 2007 Proteomic identification of proteins associated with the osmoregulatory transcription factor TonEBP/OREBP: functional effects of Hsp90 and PARP-1. Am J Physiol Renal Physiol 292 F981–F992

Chi B, O’Connell JD, Yamazaki T, Gangopadhyay J, Gygi SP, et al. 2018 Interactome analyses revealed that the U1 snRNP machinery overlaps extensively with the RNAP II machinery and contains multiple ALS/SMA-causative proteins. Scientific Reports 8 8755

Chia W, Somers WG, Wang H 2008 Drosophila neuroblast asymmetric divisions: cell cycle regulators, asymmetric protein localization, and tumorigenesis. J. Cell Biology 180 267–272

Chiang A, Priya R, Ramaswami M, Vijayraghavan K, Rodrigues V 2009 Neuronal activity and Wnt signaling act through Gsk3-β to regulate axonal integrity in mature Drosophila olfactory sensory neurons. Development 136 1273–1282

Ching W, Hang HC, Nusse R 2008 Lipid-independent secretion of a Drosophila Wnt protein. J. Biol. Chem.

Cohen ED, Tian Y, Morrisey EE 2008 Wnt signaling: an essential regulator of cardiovascular differentiation, morphogenesis and progenitor self-renewal. Development 135 789–798

Coyne AN, Yamada SB, Siddegowda BB, Estes PS, Zaepfel BL, et al. 2015 Fragile X protein mitigates TDP-43 toxicity by remodeling RNA granules and restoring translation. Human Molecular Genetics 24 6886–6898

Creugny A, Fender A, Pfeffer S 2018 Regulation of primary micro RNA processing. FEBS Lett. 592: 1980– 1996

Cutforth T, Rubin GM 1994 Mutations in Hsp83 and cdc37 impair signaling by the sevenless receptor tyrosine kinase in Drosophila. Cell 77 1027–1036

de Cárcer G, do Carmo Avides M, Lallena MJ, Glover DM, González C 2001 Requirement of Hsp90 for centrosomal function reflects its regulation of Polo kinase stability. EMBO J. 20 2878–2884

DeFranco DB, Csermely P 2000 Steroid receptor and molecular chaperone encounters in the nucleus. Science Signaling 2000 pe1-pe1

Dittmar RL, Sen S 2018 MicroRNAs in exosomes in cancer. in Cancer and Noncoding RNAs (Chakrabarti J, Mitra S Ed) Academic Press pp 59-78

Estes PS, O’Shea M, Clasen S, Zarnescu DC 2008 Fragile X protein controls the efficacy of mRNA transport in Drosophila neurons. Molecular Cellular Neuroscience 39 170–179

Farkas R, Mechler B 2000 The timing of Drosophila salivary gland apoptosis displays an l (2) gl-dose response. Cell Death Differentiation 7 89–101

Ford L, Wright W, Shay J 2002 A model for heterogeneous nuclear ribonucleoproteins in telomere and telomerase regulation. Oncogene 21 580–583

Gangaraju VK, Yin H, Weiner MM, Wang J, Huang XA, et al. 2011 Drosophila Piwi functions in Hsp90-mediated suppression of phenotypic variation. Nature genetics 43 153

Gateff E, Mechler B 1989 Tumor-suppressor genes of Drosophila melanogaster. Critical reviews in oncogenesis 1 221–245

Georgieva D, Petrova M, Molle E, Daskalovska I, Genova G 2012 Drosophila DFMR1 interacts with genes of the Lgl-pathway in the brain synaptic architecture. Biotechnology Biotechnological Equip 26 52–59

Grammatikakis N, Lin JH, Grammatikakis A, Tsichlis PN, Cochran BH 1999 p50(cdc37) acting in concert with Hsp90 is required for Raf-1 function. Mol Cell Biol 19 1661–1672

Grandbarbe L, Bouissac J, Rand M, de Angelis MH, Artavanis-Tsakonas S, et al. 2003 Delta-Notch signaling controls the generation of neurons/glia from neural stem cells in a stepwise process. Development 130 1391–1402

Guemez-Gamboa A, Coufal NG, Gleeson JG 2014 Primary cilia in the developing and mature brain. Neuron 82 511–521

Hämmerle B, Tejedor FJ 2007 A novel function of DELTA-NOTCH signalling mediates the transition from proliferation to neurogenesis in neural progenitor cells. PloS One 2 e1169

Han SP, Tang YH, Smith R 2010 Functional diversity of the hnRNPs: past, present and perspectives. Biochemical Journal 430 379–392

Hariharan IK, Bilder D 2006 Regulation of imaginal disc growth by tumor-suppressor genes in Drosophila. Annu. Rev. Genet. 40 335–361

Herr P, Hausmann G, Basler K 2012 WNT secretion and signalling in human disease. Trends Molecular Medicine 18 483–493

Hoang B, Chiba A 1998 Genetic analysis on the role of integrin during axon guidance in Drosophila. J. Neuroscience 18 7847–7855

Homem CC, Knoblich JA 2012 Drosophila neuroblasts: a model for stem cell biology. Development 139 4297–4310

Huang DW, Sherman BT, Zheng X, Yang J, Imamichi T, et al. 2009 Extracting biological meaning from large gene lists with DAVID. Current Protocols Bioinformatics 13.11. 11–13.11. 13

Humbert P, Grzeschik N, Brumby A, Galea R, Elsum I, et al. 2008 Control of tumourigenesis by the Scribble/Dlg/Lgl polarity module. Oncogene 27 6888

Humbert P, Russell S, Richardson H 2003 Dlg, Scribble and Lgl in cell polarity, cell proliferation and cancer. Bioessays 25 542–553

Inestrosa NC, Varela-Nallar L 2014 Wnt signaling in the nervous system and in Alzheimer’s disease. Jour. Mol. Cell Biology 6 64–74

Iwasaki S, Sasaki HM, Sakaguchi Y, Suzuki T, Tadakuma H, et al. 2015 Defining fundamental steps in the assembly of the Drosophila RNAi enzyme complex. Nature 521 533

Jana SC, Bettencourt-Dias M, Durand B, Megraw TL 2016 Drosophila melanogaster as a model for basal body research. Cilia 5 22

Ji Y, Tulin A 2009 Poly(ADP-ribosyl)ation of heterogeneous nuclear ribonucleoproteins modulates splicing. Nucleic Acids Research 37 3501–3513

Ji Y, Tulin A 2013 Post-transcriptional regulation by poly (ADP-ribosyl) ation of the RNA-binding proteins. International J Molecular Sciences 14 16168–16183

Jinwal UK, Abisambra JF, Zhang J, Dharia S, O’Leary JC, et al. 2012 Cdc37/Hsp90 protein complex disruption triggers an autophagic clearance cascade for TDP-43 protein. J Biol Chem 287 24814–24820

Johnson TK, Cockerell FE, McKechnie SW 2011 Transcripts from the Drosophila heat-shock gene hsr-omega influence rates of protein synthesis but hardly affect resistance to heat knockdown. Molecular genetics and genomics 285 313–323

Karak S, Jacobs JS, Kittelmann M, Spalthoff C, Katana R, et al. 2015 Diverse roles of axonemal dyneins in Drosophila auditory neuron function and mechanical amplification in hearing. Scientific Reports 5 17085

Kassis JA, Kennison JA, Tamkun JW 2017 Polycomb and trithorax group genes in Drosophila. Genetics 206 1699–1725

Kato K, Forero MG, Fenton JC, Hidalgo A 2011 The glial regenerative response to central nervous system injury is enabled by pros-notch and pros-NF?B feedback. PLoS Biology 9 e1001133

Klämbt C, Schmidt O 1986 Developmental expression and tissue distribution of the lethal (2) giant larvae protein of Drosophila melanogaster. EMBO J. 5 2955–2961

Korochkina L, Fursenko O, Sherudilo A 1975 Characteristics of the endocrine system in Drosophila melanogaster 1 (2) gl mutants, differing in time of death. Genetika 11 57–65

Lackie RE, Maciejewski A, Ostapchenko VG, Marques-Lopes J, Choy W-Y, et al. 2017 The Hsp70/Hsp90 chaperone machinery in neurodegenerative diseases. Frontiers Neuroscience 11 254

Lakhotia SC 2011 Forty years of the 93D puff of Drosophila melanogaster. J. Biosciences 36 399–423

Lakhotia SC 2016 Non-coding RNAs have key roles in cell regulation. Proc. Indian Natn. Sci. Acad. 82 in press

Lakhotia SC 2017 From heterochromatin to long noncoding RNAs in Drosophila: Expanding the arena of gene function and regulation. in Long Non Coding RNA Biology (Rao M R S Ed) Springer Nature Singapore Pte Ltd pp 75-118

Lakhotia SC, Mallik M, Singh AK, Ray M 2012 The large noncoding hsr?-n transcripts are essential for thermotolerance and remobilization of hnRNPs, HP1 and RNA polymerase II during recovery from heat shock in Drosophila. Chromosoma 121 49–70

Lakhotia SC, Ray P 1996 hsp 83 mutation is a dominant enhancer of lethality associated with absence of the non–protein coding hsr omega locus in Drosophila melanogaster. Journal of Biosciences 21 207– 219

Lange BM, Bachi A, Wilm M, González C 2000 Hsp90 is a core centrosomal component and is required at different stages of the centrosome cycle in Drosophila and vertebrates. EMBO J. 19 1252–1262

Li B, Carey M, Workman JL 2007 The role of chromatin during transcription. Cell 128 707-719

Lim J, Norga KK, Chen Z, Choi KW 2005 Control of planar cell polarity by interaction of DWnt4 and four-jointed. genesis 42 150–161

Lin H, Schagat T 1997 Neuroblasts: a model for the asymmetric division of stem cells. Trends Genetics 13 33–39

Louvi A, Grove EA 2011 Cilia in the CNS: the quiet organelle claims center stage. Neuron 69 1046-1060

Mallik M, Lakhotia SC 2009 RNAi for the large non-coding hsr omega transcripts suppresses polyglutamine pathogenesis in Drosophila models. Rna Biology 6 464–478

Mallik M, Lakhotia SC 2011 Pleiotropic consequences of misexpression of the developmentally active and stress-inducible non-coding hsr? gene in Drosophila. J. Biosciences 36 265–280

Martinek N, Shahab J, Saathoff M, Ringuette M 2008 Haemocyte-derived SPARC is required for collagen-IV-dependent stability of basal laminae in Drosophila embryos. J. Cell science 121 1671–1680

Mazaira GI, Camisay MF, De Leo S, Erlejman AG, Galigniana MD 2016 Biological relevance of Hsp90-binding immunophilins in cancer development and treatment. Int. J. Cancer 138 797–808

McGurk L, Gomes E, Guo L, Mojsilovic-Petrovic J, Tran V, et al. 2018 Poly (ADP-Ribose) prevents pathological phase separation of TDP-43 by promoting liquid demixing and stress granule localization. Molecular Cell 10.1016/j.molcel.2018.07.002 10.1016/j.molcel.2018.1007.1002

Merz R, Schmidt M, Török I, Protin U, Schuler G, et al. 1990 Molecular action of the l (2) gl tumor suppressor gene of Drosophila melanogaster. Environmental Health Perspectives 88 163

Metsalu T, Vilo J 2015 ClustVis: a web tool for visualizing clustering of multivariate data using Principal Component Analysis and heatmap. Nucleic Acids Research 43 W566–W570

Meyer S, Schmidt I, Klämbt C 2014 Glia ECM interactions are required to shape the Drosophila nervous system. Mech. Development 133 105–116

Miyoshi T, Takeuchi A, Siomi H, Siomi MC 2010 A direct role for Hsp90 in pre-RISC formation in Drosophila. Nature Structural Molecular Biology 17 1024

Moore DJ, Onoufriadis A, Shoemark A, Simpson MA, Zur Lage PI, et al. 2013 Mutations in ZMYND10, a gene essential for proper axonemal assembly of inner and outer dynein arms in humans and flies, cause primary ciliary dyskinesia. Amer.J. Human Genetics 93 346–356

Morcillo G, Diez JL, Carbajal ME, Tanguay RM 1993 HSP90 associates with specific heat shock puffs (hsr?) in polytene chromosomes of Drosophila and Chironomus. Chromosoma 102 648–659

Ohshiro T, Yagami T, Zhang C, Matsuzaki F 2000 Role of cortical tumour-suppressor proteins in asymmetric division of Drosophila neuroblast. Nature 408 593

Oliva CA, Vargas JY, Inestrosa NC 2013 Wnts in adult brain: from synaptic plasticity to cognitive deficiencies. Front. Cellular Neuroscience 7 224

Olivieri D, Senti K-A, Subramanian S, Sachidanandam R, Brennecke J 2012 The cochaperone shutdown defines a group of biogenesis factors essential for all piRNA populations in Drosophila. Molecular Cell 47 954–969

Onorati MC, Lazzaro S, Mallik M, Ingrassia AM, Carreca AP, et al. 2011 The ISWI chromatin remodeler organizes the hsr? ncrna–containing omega speckle nuclear compartments. PLoS Genetics 7 e1002096

Pandey R, Blanco J, Udolph G 2011 The glucuronyltransferase GlcAT-P is required for stretch growth of peripheral nerves in Drosophila. Plos One 6 e28106

Patel-King RS, King SM 2016 A prefoldin-associated WD-repeat protein (WDR92) is required for the correct architectural assembly of motile cilia. Molecular Biology Cell 27 1204–1209

Piccolo LL, Bonaccorso R, Attardi A, Li Greci L, Romano G, et al. 2018 Loss of ISWI Function in Drosophila nuclear bodies drives cytoplasmic redistribution of Drosophila TDP-43. Int. J. Mol. Sci. 19 1082

Piccolo LL, Corona D, Onorati MC 2014 Emerging roles for hnRNPs in post-transcriptional regulation: what can we learn from flies? Chromosoma 123 515–527

Piccolo LL, Jantrapirom S, Nagai Y, Yamaguchi M 2017 FUS toxicity is rescued by the modulation of lncRNA hsr? expression in Drosophila melanogaster. Scientific Reports 7 15660

Piccolo LL, Yamaguchi M 2017 RNAi of arcRNA hsr? affects sub-cellular localization of Drosophila FUS to drive neurodiseases. Experimental Neurology 292 125–134

Prasanth K, Rajendra T, Lal A, Lakhotia S 2000 Omega speckles-a novel class of nuclear speckles containing hnRNPs associated with noncoding hsr-omega RNA in Drosophila. Journal of cell science 113 3485–3497

Pratt WB, Toft DO 1997 Steroid receptor interactions with heat shock protein and immunophilin chaperones. Endocr Rev 18 306–360

Ray M, Lakhotia SC 2017 Altered hsr? lncRNA levels in activated Ras background further enhance Ras activity in Drosophila eye and induces more R7 photoreceptors. bioRxiv 10.1101/224543 224543

Reichsman F, Smith L, Cumberledge S 1996 Glycosaminoglycans can modulate extracellular localization of the wingless protein and promote signal transduction. J. Cell Biology 135 819–827

Rhee DY, Cho D-Y, Zhai B, Slattery M, Ma L, et al. 2014 Transcription factor networks in Drosophila melanogaster. Cell Reports 8 2031–2043

Ringrose L 2017 Noncoding RNAs in polycomb and trithorax regulation: A quantitative perspective. Annu Review Genetics 51 385–411

Riparbelli MG, Callaini G, Megraw TL 2012 Assembly and persistence of primary cilia in dividing Drosophila spermatocytes. Developmental Cell 23 425–432

Rogowski K, Juge F, Van Dijk J, Wloga D, Strub J-M, et al. 2009 Evolutionary divergence of enzymatic mechanisms for posttranslational polyglycylation. Cell 137 1076–1087

Romano M, Feiguin F, Buratti E 2016 TBPH/TDP-43 modulates translation of Drosophila futsch mRNA through an UG-rich sequence within its 51 UTR. Brain Research 1647 50-56

Roy S, Lakhotia S 1991 In situ patterns of nuclear replication in brain ganglia ofl (2) gl 4 mutant larvae of Drosophila melanogaster. Journal of Genetics 70 161–168

Ruden DM, Lu X 2008 Hsp90 affecting chromatin remodeling might explain transgenerational epigenetic inheritance in Drosophila. Current Genomics 9 500–508

Rutherford S, Hirate Y, Swalla BJ 2007 The Hsp90 capacitor, developmental remodeling, and evolution: the robustness of gene networks and the curious evolvability of metamorphosis. Critical Reviews Biochemistry Molecular Biology 42 355–372

Rutherford SL, Lindquist S 1998 Hsp90 as a capacitor for morphological evolution. Nature 396 336

Sangster TA, Queitsch C, Lindquist S 2003 Hsp90 and chromatin: where is the link? Cell Cycle 2 165-167 Satir P,

Pedersen LB, Christensen ST 2010 The primary cilium at a glance. J Cell Sci 123 499–503

Sato M 2006 Upregulation of the Wnt/β-catenin pathway induced by transforming growth factor-β in hypertrophic scars and keloids. Acta Dermato-Venereologica 86 300–307

Sawarkar R, Sievers C, Paro R 2012 Hsp90 globally targets paused RNA polymerase to regulate gene expression in response to environmental stimuli. Cell 149 807–818

Schmidt I, Thomas S, Kain P, Risse B, Naffin E, et al. 2012 Kinesin heavy chain function in Drosophila glial cells controls neuronal activity. Journal of Neuroscience 32 7466–7476

Schuettengruber B, Bourbon H-M, Di Croce L, Cavalli G 2017 Genome regulation by polycomb and trithorax: 70 years and counting. Cell 171 34–57

Segnitz B, Gehring U 1997 The function of steroid hormone receptors is inhibited by the hsp90-specific compound geldanamycin. Journal of Biological Chemistry 272 18694–18701

Singh AK, Lakhotia SC 2015 Dynamics of hnRNPs and omega speckles in normal and heat shocked live cell nuclei of Drosophila melanogaster. Chromosoma 124 367–383

Specchia V, D’Attis S, Puricella A, Bozzetti M 2017 dFmr1 Plays Roles in Small RNA Pathways of Drosophila melanogaster. Int. J. Mol. Sci. 18 1066

Specchia V, Piacentini L, Tritto P, Fanti L, D’Alessandro R, et al. 2010 Hsp90 prevents phenotypic variation by suppressing the mutagenic activity of transposons. Nature 463 662–665

Tapadia M, Lakhotia S 1997 Specific induction of the hsr? locus of Drosophila melanogaster by amides. Chromosome Research 5 359–362

Tariq M, Nussbaumer U, Chen Y, Beisel C, Paro R 2009 Trithorax requires Hsp90 for maintenance of active chromatin at sites of gene expression. Proceedings of the National Academy of Sciences pnas. 0809669106

Trapnell C, Roberts A, Goff L, Pertea G, Kim D, et al. 2012 Differential gene and transcript expression analysis of RNA-seq experiments with TopHat and Cufflinks. Nature protocols 7 562–578

van der Straten A, Rommel C, Dickson B, Hafen E 1997 The heat shock protein 83 (Hsp83) is required for Raf-mediated signalling in Drosophila. EMBO J. 16 1961–1969

Welch RM 1957 A developmental analysis of the lethal mutant l (2) gl of Drosophila melanogaster based on cytophotometric determination of nuclear desoxyribonucleic acid (DNA) content. Genetics 42 544

Xie X, Auld VJ 2011 Integrins are necessary for the development and maintenance of the glial layers in the Drosophila peripheral nerve. Development 138 3813–3822

Youn J-Y, Dunham WH, Hong SJ, Knight JD, Bashkurov M, et al. 2018 High-density proximity mapping reveals the subcellular organization of mRNA-associated granules and bodies. Molecular Cell 69 517- 532. e511

Zarnescu DC, Jin P, Betschinger J, Nakamoto M, Wang Y, et al. 2005 Fragile X Protein Functions with Lgl and the PAR Complex in Flies and Mice. Developmental Cell 8 43–52

Zhang L, Chen Q, An W, Yang F, Maguire EM, et al. 2017 Novel pathological role of hnRNPA1 (heterogeneous nuclear ribonucleoprotein A1) in vascular smooth muscle cell function and neointima hyperplasia. Arteriosclerosis, Thrombosis Vascular Biology 37 2182–2194

Zhang Q-S, Manche L, Xu R-M, Krainer AR 2006 hnRNP A1 associates with telomere ends and stimulates telomerase activity. Rna 12 1116–1128

Zhao R, Kakihara Y, Gribun A, Huen J, Yang G, et al. 2008 Molecular chaperone Hsp90 stabilizes Pih1/Nop17 to maintain R2TP complex activity that regulates snoRNA accumulation. J Cell Biol 180 563–578

